# The Gut Microbiome Attenuates the Adaptation of a Reproductive Endosymbiont

**DOI:** 10.1101/2024.09.10.611936

**Authors:** Madangchanok Imchen, Rupinder Kaur, Sebastian Leon-Fallas, Darlene Huilin Zhou Luo, Mia W. Bitman, Emilie Lefoulon, Angelina McGarry, Leah R. Carpenter, Cheyenne Lehman, Sarah R. Bordenstein, Seth R. Bordenstein

**Affiliations:** Departments of Biology and Entomology, One Heath Microbiome Center, Huck Institutes of the Life Sciences, The Pennsylvania State University, USA; Department of Biological Sciences, University of Illinois Chicago, IL, USA

**Keywords:** cytoplasmic incompatibility, microbiome, intracellular bacteria, endosymbiosis, *Drosophila melanogaster*

## Abstract

Holobionts contain spatially separated microbes whose interorgan effects are a neglected source of potential, functional variation. Here we show the extracellular gut microbiome modulates the transcriptome and adaptation of the endosymbiont *Wolbachia* in *Drosophila melanogaster* testes. Gut bacteria suppress cytoplasmic incompatibility (CI), a sperm-egg lethality trait, by altering endosymbiont pathogenicity gene expression. Transgenic assays identify the *enchancer of cytoplasmic incompatibility* gene *eci* as causal to the variation. Restoring the gut microbiome in laboratory and wild-caught, germ-free flies recapitulates CI attenuation and shifts host fitness. Intact holobionts sustain higher densities of the two symbiont types versus isolated rearing, revealing a synergistic, replication interaction between the symbiont classes. The ‘gut-germline’ axis on symbiont functions has far-reaching implications for the study of holobiont trait variation.

## Main

Holobionts harbor layered microbial consortia - from obligate intracellular endosymbionts to extracellular communities(*1–3*) - that can hypothetically engage in competitive, collaborative, or stochastic interplay(*4*). Such cross-compartment interactions may trigger cascading effects on holobiont biology, yet they remain fundamentally understudied because conventional antibiotic regimes are indiscriminate and endosymbiont cultivation remains challenging(*5*). Here, we circumvent these limitations using a tractable, model system that uncouples a vertically-inherited reproductive endosymbiont from an independently manipulable, environmentally-acquired, gut microbiome.

In the *Drosophila melanogaster* holobiont, the α-proteobacterium *Wolbachia* is a canonical reproductive endosymbiont and one of the most ecologically-pervasive, maternally-inherited bacteria in animals(*6*). Within the reproductive tract, *Wolbachia* catalyze cytoplasmic incompatibility (CI), a paternal-effect, epigenetic lethality where crosses between symbiotic males and aposymbiotic females result in embryonic and larval death(*7–12*). Lethality is rescued when females harbor the same *Wolbachia* strain, selfishly driving *Wolbachia’s* rapid spread(*8*). In parallel, *D. melanogaster* maintains a simple gut microbiome of a few core taxa, easily removed to generate germ-free hosts(*13*). This tractability – altering and restoring symbiont classes independently or combined – makes *D. melanogaster* suited for dissecting interorgan symbiotic dynamics.

While *Wolbachia* are broadly detectable in somatic tissues(*14*), they are excluded from the *D. melanogaster* gut lumen, residing within epithelial cells without inducing immune responses(*15*). This spatial architecture precludes direct antagonism or synergy with the luminal gut microbiome. Nevertheless, indirect associations occur where *Wolbachia* presence or absence associates with gut microbiome shifts, though antibiotic exposure may be the primary driver(*15*, *16*). Moreover, midgut-testes physical adjacency(*17*) enables a plausible interaction between anatomically-separated bacteria. Thus, there is a critical gap of whether trade-offs and synergies occur locally or systematically between distinct symbiont classes across anatomical compartments. Past work largely focused on binary symbioses or single anatomical sites, leaving emergent interactions across a symbiotic system understudied. We address three questions to unravel interorgan, symbiotic functions: (i) Can a reproductive endosymbiont (*Wolbachia*) function autonomously from the gut microbiome? (ii) What effects does the gut microbiome exert on endosymbiont densities, gene expression, and major adaptation? And (iii) Do the two symbiotic classes independently and/or synergistically shape each other’s fitness and that of the host? We uncover a previously, overlooked impact of spatially- and lifestyle-distinct bacteria on holobiont-level traits.

## Results

We generated *D. melanogaster* with and without *Wolbachia* and the gut microbiome by dechorionating the fertilized eggshell layer, eliminating the extracellular gut microbiome(*13*) while preserving the transovarially-inherited, reproductive endosymbiont. We adhere to the typical nomenclature of conventional (CV) and germ-free (GF) hosts with *Wolbachia* (CV+ and GF+) and without it (CV- and GF-) (Fig 1A). We also generated germ-free hosts with a CV+ fecal microbiome transplant (GF_M_+) to assess causal recapitulation of microbiome effects. We confirmed *Wolbachia* infection by PCR, GF+/- status by culturing whole fly homogenates (see methods) (Fig S1), and that the CV+/- lines share the same genetic makeup (Table S1).

**Fig 1.**
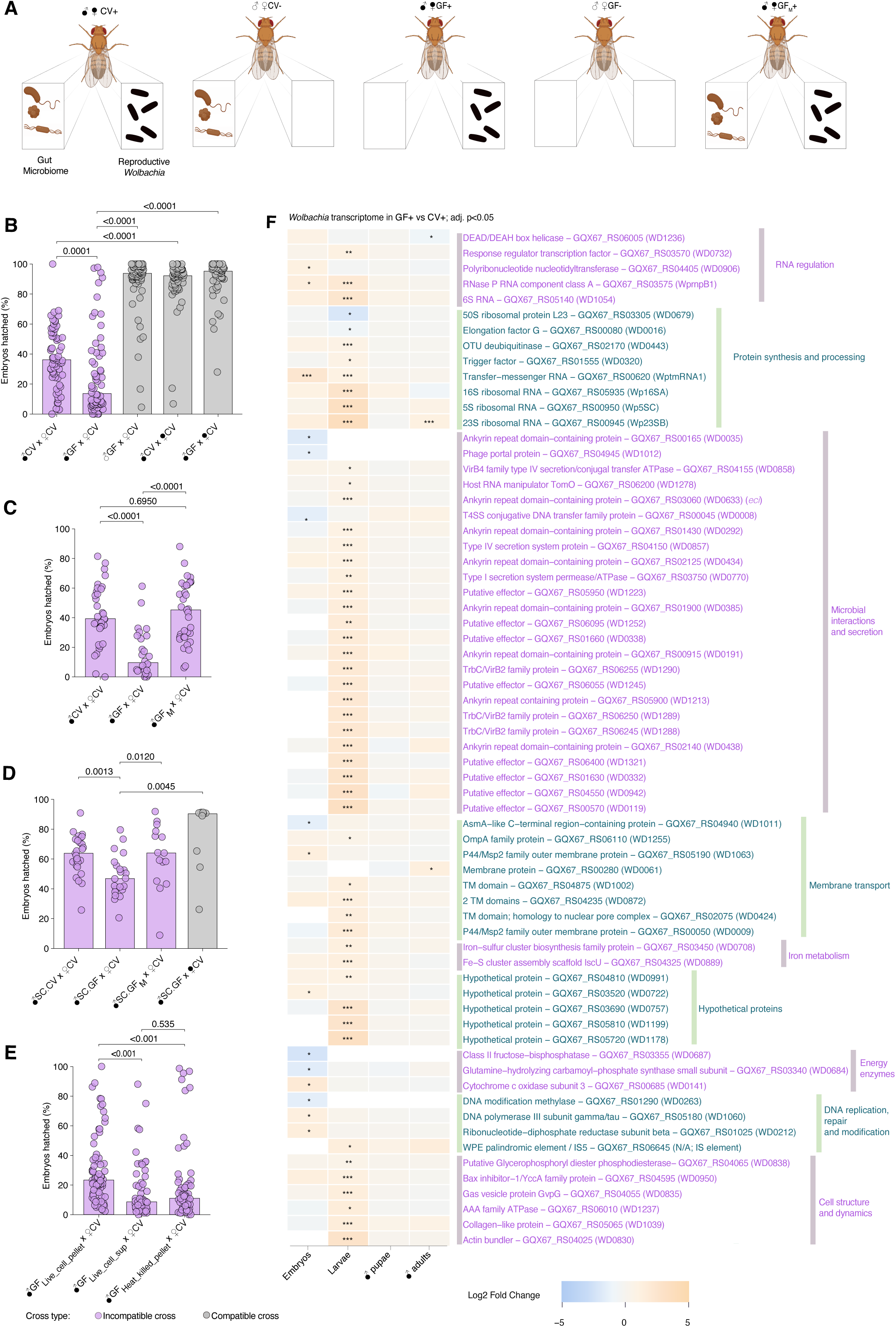
The gut microbiome attenuates CI penetrance and *Wolbachia* symbiosis transcripts. (A) Schematic representation and notation of the fly lines showing (from left to right): complete symbiont community, gut microbiome only, endosymbiont (*Wolbachia*) only, symbiont-free, and fecal microbiome reintroduced into a endosymbiont (*Wolbachia*) only host. (B) Germ-free males with *Wolbachia* (GF+) induce significantly higher CI (dark pink, measured as the percentage of hatching embryos into larvae) than conventional (CV+) males. Compatible, control crosses (grey bars) with normal embryonic hatching percentages express either no CI or rescue of CI, as expected. (C) GF+ males with a reintroduced, native microbiome (GF+_M_) recapitulate the attenuated CI similar to CV+ males. (D) Natural fly lines from State College (SC), Pennsylvania confirm that the gut microbiome causally attenuates the CI phenotype in flies from SC.CV natural populations and SC.GF+_M_ males receiving a fecal microbiome reintroduction. (E) The attentuaion of CI requires a live cell pellet with viable, gut bacterial cells relative to its purified supernatant or heatkilled gut bacterial pellet. Each data point in panels B-E represents a single pair mating. Vertical bars represent the median, and P-values are based on a Kruskal-Wallis test followed by Benjamini-Hochberg false discovery rate (FDR) multiple correction test. (F) Comparative *Wolbachia* transcriptomes across development reveal the number and types of *Wolbachia* genes that signficantly and differentially expressed between GF+ and CV+, with marked enrichment in the larval stage when germcell development commences. Labels on the y-axis are based on the GQX67 locus tags (NZ_CP046925.1), with the corresponding WD locus tags (AE017196) shown in parentheses.

### Live gut microbiome causally attenuates the adaptation of a reproductive endosymbiont

Reproductive *Wolbachia* and gut bacteria of *D. melanogaster* have largely been studied independently(*18*, *19*). To address their interorgan connectivity, we evaluated if the intestinal microbiome shapes *Wolbachia*’s defining CI phenotype. We first assess CI level variation with and without the gut microbiome using pairwise crosses between virgin males with *Wolbachia* (CV+ and GF+) and aposymbiotic females lacking *Wolbachia* (CV-). The CI paternal-effect lethality was scored using our traditional assays as the percent reduction of embryonic hatching into larvae. Both CV+ and GF+ males induce CI, confirming *Wolbachia* express the trait autonomously regardless of gut microbiome status. However, GF+ induce a 2.6-fold stronger CI (median embryonic hatch rate = 13.66%) relative to CV+ males (median = 36.17%) (Fig 1B). This amplified lethality was rescuable by CV+ females, confirming bona fide CI rather than an artifactual host sterility. Restoring the native microbiome via fecal-oral transmission notably reduces CI attenuation (Fig 1C, median = 45.28%), establishing the gut microbiome as a causal modulator of *Wolbachia*-induced lethality.

Consistent with the laboratory strains, microbiome elimination in wild-caught males flies from State College, Pennsylvania, USA, denoted SC.GF+, significantly strengthens CI (median = 46.89%) relative to their conventional counterparts, SC.CV+ flies (median = 63.83%) (Fig 1D). Crucially, native microbiome transplantation restored CI to conventional levels (median = 64.07%), confirming this modulation is not an artifact of prolonged laboratory culture. Field-derived *D. melanogaster* characteristically express weaker CI than laboratory strains(*20*), yet CI here remains statistically rescuable to a fully compatible hatch rate of 90.36%.

We next tested whether CI attenuation via the gut microbiome requires live microbial cells. The *D. melanogaster* gut microbiome is predominantly composed of *Lactobacillaceae* and *Acetobacteraceae*(*13*). Only live cells attenuated CI; 0.20 µm-filtered supernatant and heat-killed pellet failed to phenocopy this suppression (Fig 1E). The partial attenuation of CI via the supernatant likely reflects the limitations of a single-timepoint delivery that cannot simulate continuous bacterial growth(*21*). Together, gut microbiome causally suppresses *Wolbachia*-induced CI in laboratory and natural populations, requiring live microbial cells.

### The gut microbiome primes virulence gene expression of the reproductive endosymbiont

Having established that the gut microbiome causally modulates the endosymbiont adaptation, we evaluated effects on *Wolbachia* gene expression across host developmental stages. Transcriptional activity of symbiotic bacteria alter phenotypes within specific organs(*22–25*), yet its inter-organ impacts on reproductive symbiont gene expression is unknown. To probe this interorgan axis, we profiled *Wolbachia* transcriptomes across embryos, larvae, male pupae, and male adults in GF+ and CV+ flies, interrogating if gut microbiome development bestows a molecular signature.

Among the stages examined, the larval transcriptome most differentiates GF+ from CV+ (Figure 1F). The 51 significant differentially-expressed genes in larvae showed an average increase of 3.91-fold spanning 0.23-fold underexpressed to 12.69-fold overexpressed. The larval specificity is noteworthy, marking the commencement of gut and testes development and thus when they are most sensitive to perturbation. During this stage, their body mass(*26*) and intestinal bacteria(*27*) increase by two orders of magnitude, while spermatogonia and spermatocytes store RNA transcripts and regulatory non-coding RNAs for spermatid remodeling downstream that impact CI(*28*).

Most altered *Wolbachia* transcripts map to Microbial Interactions and Secretion, encompassing hyperexpression of type IV secretion system (T4SS) genes, virulence effectors, and ankyrin-containing proteins widely involved in host-microbe relationships (Fig 1F). They are also differentially expressed in larvae across multiple *Wolbachia* transcriptomic studies(*29–31*). Thus, gut microbiome absence amplifies active *Wolbachia* transcriptional components, suggesting a tunable sensitivity of host interaction genes. Changes in protein synthesis genes and membrane transport further suggest *Wolbachia* shift to a state of advancing host interactions in GF+ larvae.

An intriguing candidate upregulated in GF+ flies was GQX67_RS03060 (WD0633), an ankyrin repeat domain–containing protein (GF+ vs. CV+ log2FC = 0.772, p < 0.001) (Fig 1F). We named this gene *eci* for *enhancer of cytoplasmic incompatibility.* We focused on this candidate because (i) *eci* is positioned in the *Wovirus* region harboring the eukaryotic association module (ii) it is immediately downstream of the *cytoplasmic incompatibility factor* genes *cifA* and *cifB*(*32, 33*) (iii) it contains an OTU deubiquitinase domain and multiple ankyrin repeats implicated in host-microbe interactions (Fig 2A), (iv) it contains sequence motifs predictive of T4SS effectors(*34*, *35*) and was the top effector prediction in *w*Mel(*36*). Previous transgenic assays showed *eci* alone does not induce a CI(*37*),(*35*); and GF+ larvae/adult males show neither higher *Wolbachia* densities nor increased *cif* expression compared to CV+ counterparts (see below, Fig 4C-F). This suggests that CI in GF+ hosts may be enhanced by effector proteins independent of other common factors.

**Fig 2.**
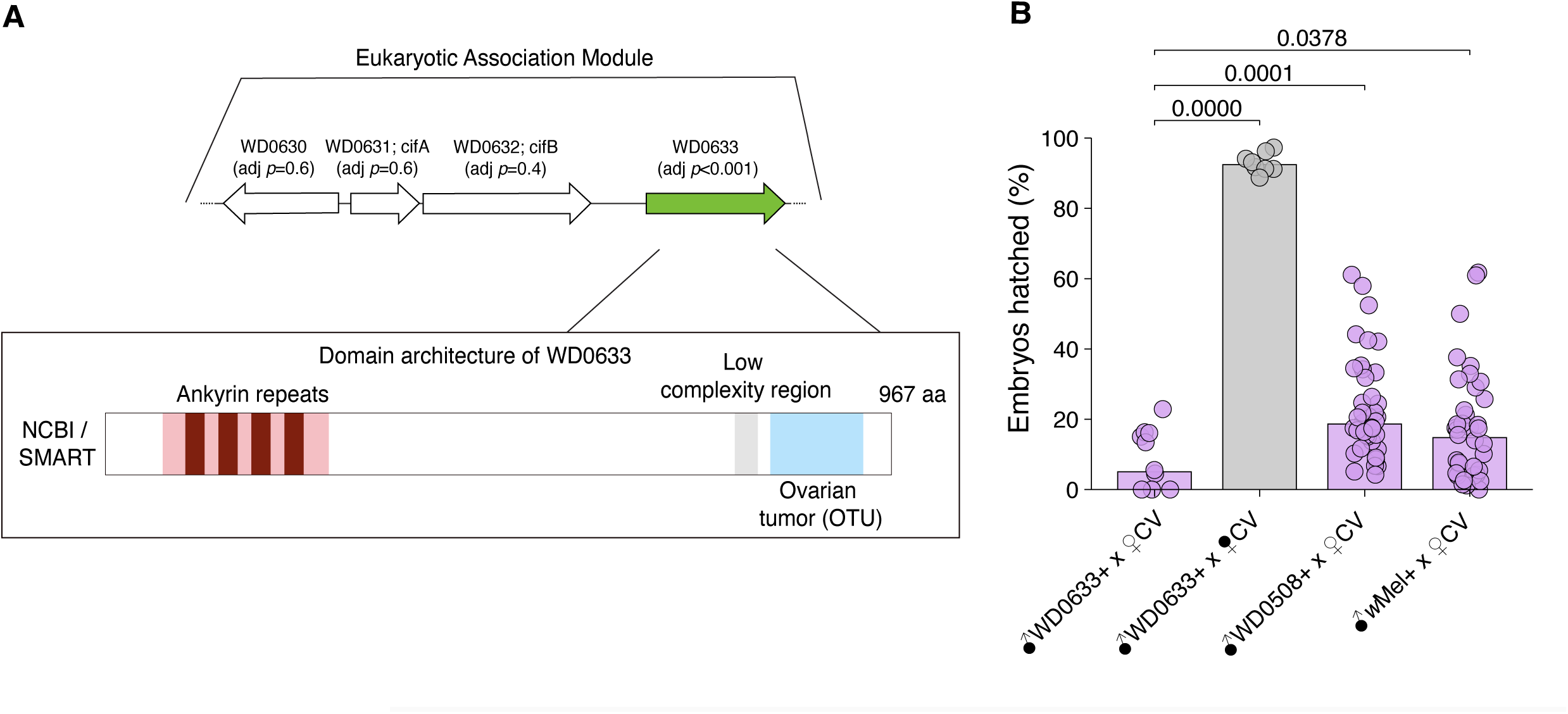
Overexpression of the *eci* gene enhances cytoplasmic incompatibility. (A) The *enhancer of cytoplasmic incompatibility* gene, *eci* (WD0633), resides 3′ downstream of *cifB*. It is a 967-amino-acid protein composed of two main domains: ankyrin repeats and an ovarian tumor (OTU) deubiquitinase domain. (B) Overexpression of *eci* increased wild-type (*w*Mel+) CI compared to overexpression of the control gene WD0633 and natural wild-type CI, which was rescued by crossing with *Wolbachia*-infected females.

To test whether *eci* augments CI penetrance in GF+ males, we utilized the GAL4/UAS system to overexpress the codon-optimized transgene in CV+ testes. *eci* expression strengthened CI, as predicted, in crosses to uninfected females, an effect rescued by infected females, as expected (Fig 2B). Collectively, these findings demonstrate that the gut microbiome restrains *Wolbachia* virulence gene expression, with *eci* serving as a key effector whose microbiome-dependent expression level scales with CI intensity.

### The gut microbiome tunes gene expression and cell biological features of sperm chromatin remodeling

Since CI causes histone retention and protamine depletion in developing sperm(*7*, *38*) (Fig 3A), we hypothesized that the increases in *Wolbachia* gene expression manifest in regulatory shifts between GF+ and CV+ late male pupae, when this nucleoprotein exchange begins in *D. melanogaster* male gametes(*39*, *40*). For histone biology, the significant effect that differentiates GF+ (vs. to CV+) from GF- (vs. to CV-) is *His3.3B* (Fig S2) whose expression exhibits a 1.84-fold increase (Table S2, Log2FC= 0.88, p < 0.05). H3.3 histone is removed in the early canoe stage during the protein transition(*41*). CI specifically causes a defect in H3.3/H4 complex arrangement on the paternal chromatin after fertilization(*11*). Furthermore, other core nucleosome components, *His2Av*, *His2B*, *His3.3A*, *His3.3B*, and *His4r* (*42*), show a non-significant upregulation trend in GF+ pupae (Fig S2). These results link to CI-defining histone retention documented in flies(*7*) and mosquitoes(*38*). Consequently, *Wolbachia* transcriptional variation links with enhanced histone-related gene expression in the GF+ state.

**Fig 3.**
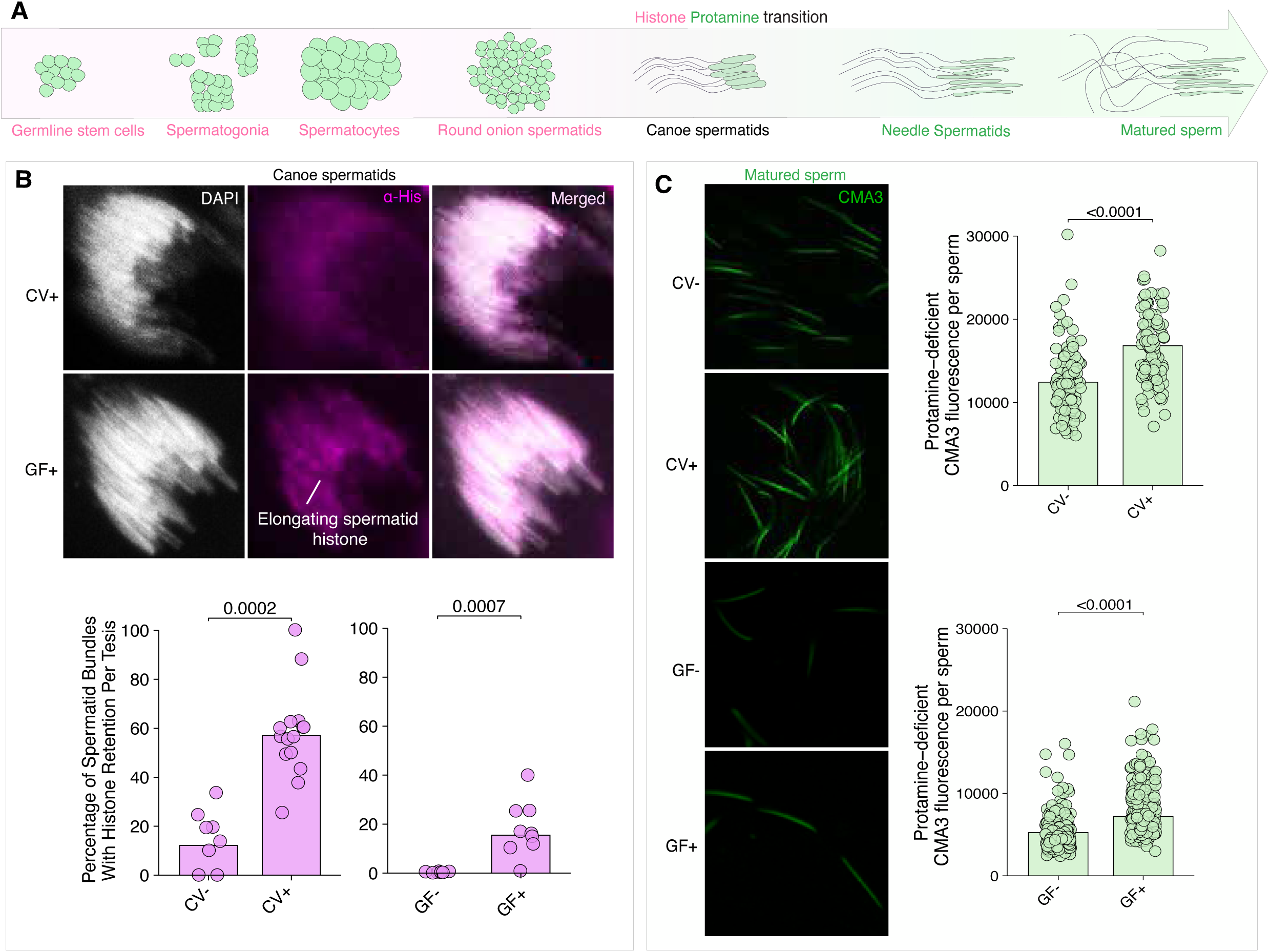
CI defects occur during the histone-to-protamine transition of sperm development. (A) During *Drosophila melanogaster* sperm development, chromatin is initially packaged with histones. In the late canoe stage, histones are normally replaced by protamines and protamine-like proteins, facilitating tight chromatin condensation within the sperm head. However, in CI testes, abnormal histone retention occurs leading to ensuring protamine deficiency in mature sperm. (B) Testes (N = ∼10) from <8-hour-old virgin males of GF-, GF+, CV-, and CV+ lines were dissected and immunostained to visualize and quantify histone abundance (magenta) in bundles of late-canoe elongating spermatids during spermiogenesis and the percentage of spermatid bundles with histone retention. DAPI stain (grey) labels spermatid nuclei. GF+ and CV+ late canoe spermatids both significantly retain histones compared to their negative counterparts GF- and CV-, respectively. (C) Fluorescent CMA3 stains DNA regions of mature sperm with protamine deficiency. Individual sperm head intensity (green) was quantified in ImageJ. GF+ and CV+ mature sperm have enhanced CMA3 intensity, consistent with protamine deficiency levels compared to GF- and CV- negative controls, respectively. In B and C, vertical bars represent the median, and p-values were calculated using the Mann-Whitney U test in R.

Four protamine and protamine-like genes are involved in mature sperm formation(*43*, *44*). Consistent with CI’s causal role in depleting protamine abundance in sperm(*7*, *38*), *ProtB*, *Prtl99C*, and *Mst77F* are significantly downregulated in GF+ vs. CV+ males (Fig S2). *ProtB* is significantly downregulated in GF+ vs. CV+ males (Table S2, fold change = 0.4, Log2FC = - 1.238, p < 0.02) but not in GF- vs. CV- males. This gene is a key structural protein for male fertility and sperm nuclear shape, both of which are disrupted in CI-causing males(*45*). Moreover, most male-specific transcript (*Mst*) genes involved in secretory proteins, protamine-based chromatin remodeling, and spermatid maturation decreased in GF+ males, including *Mst77Y-13* and *Mst84B* that are significantly reduced in symbiotic males only (Table S2, fold change = 0.36 & 0.40, Log2FC = -1.48 & -1.32 p < 0.02). Thus, *Wolbachia* generally weaken gene expression of sperm-associated proteins, some of which contribute to the protamine deficiency observed in CI sperm. Collectively, transcriptomic alterations in key CI cellular pathway genes link to augmented CI penetrance and *Wolbachia* virulence gene expression in germ-free flies.

We next leveraged cell biological markers of sperm chromatin integrity to evaluate CI-nucleoprotein signatures in GF+, GF-, CV+, and CV- testes. Using a generalized core histone antibody, histone retention occurs in both GF+ and CV+ late canoe spermatid bundles compared to GF- and CV-, respectively (Fig 3B). GF+ exhibit a median of 15.48% histone retention vs. 0% for GF-, and CV+ show 57.14% histone retention vs. 12.14% for CV-. As such, quantitative differences occur in CV testes, while qualitative differences in histone retention occur in GF+ testes.

Finally, chromomycin A3 stain binds DNA regions devoid of or loosely attached to protamines and thus determines if CI sperm suffer from a generalized, protamine defect. Cytochemical staining of mature sperm indicate protamine deficiency is 1.45-fold elevated in GF+ vs. GF- males (Fig 3C), slightly exceeding the 1.3-fold difference in CV+ vs. CV- males. Together, these results demonstrate sperm integrity changes *in situ* underpin CI, as expected. Our previous genetic work demonstrated that enhanced protamine deficiency is causal to increases in CI penetrance(*28*).

### The integrated holobiont amplifies the abundance of gut and reproductive bacteria relative to each constituent alone

CI strength varies with *Wolbachia* densities(*46*), expression of *cifA* and *cifB*(*47*), host genetics, among other factors(*48*). To test if the gut microbiome attenuates CI by lowering *Wolbachia* densities, we quantified relative *Wolbachia* amounts (*groEL)* normalized to the *D. melanogaster* housekeeping gene *rp49*. Unexpectedly, *Wolbachia* density is 2.07-fold lower (p<0.05) in strong CI-inducing GF+ vs. CV+ males, and it is equal in larvae (Fig 4A,B and S3). Thus, the intact CV+ adult holobiont has higher *Wolbachia* copies than GF+ males. Moreover, *cifA* and *cifB* gene expression is unchanged compared to the *Wolbachia* housekeeping gene *groEL* (Fig 4E,F). Thus, *cif* transcription levels are independent of the CI change, *Wolbachia* density decline, and loss of the extracellular microbiome. These findings indicate enhanced CI in GF+ males is not attributable to increased symbiont abundance or *cif* gene expression.

**Fig 4.**
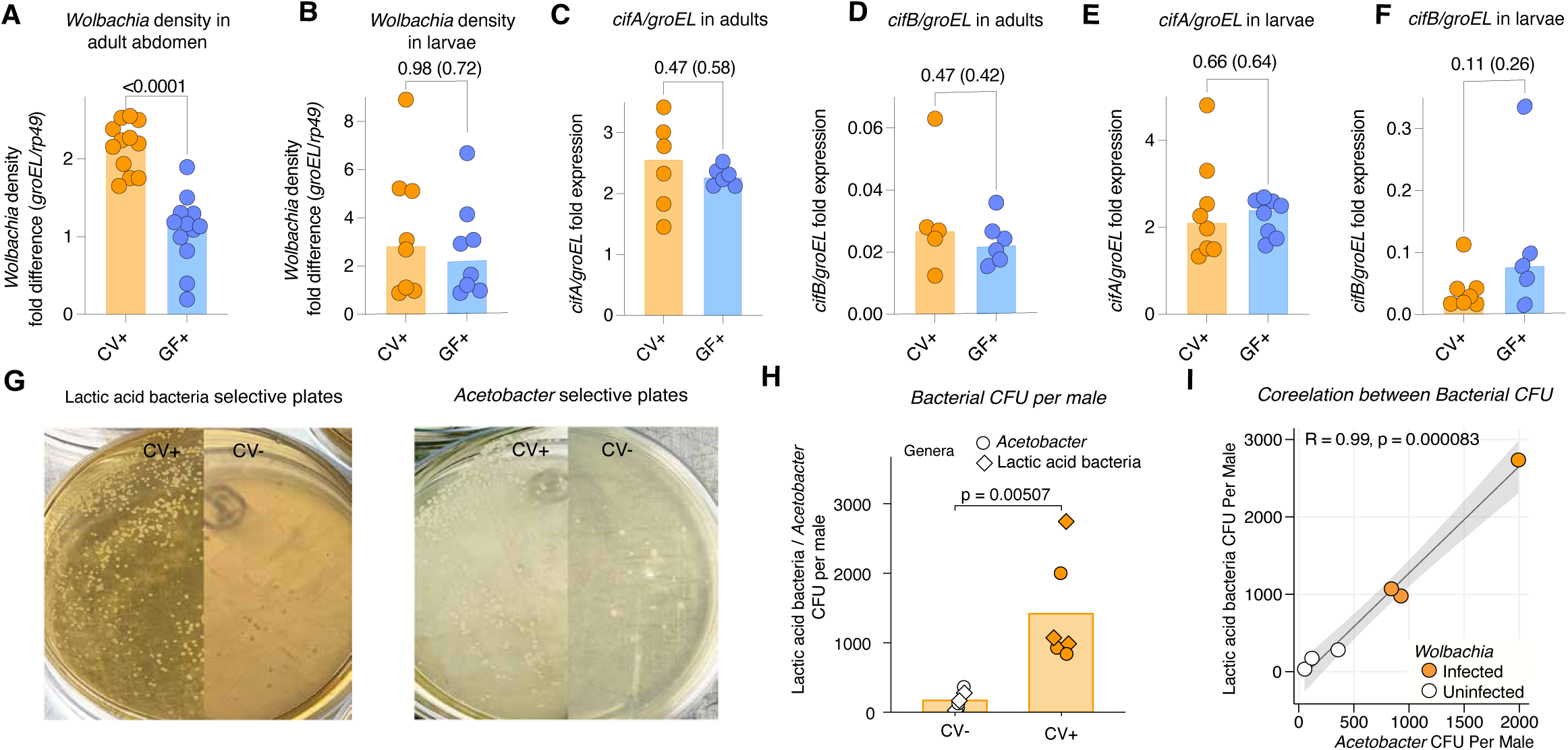
Gut microbiome proliferation in CI causing males occurs independenlty of CI gene expression. *Wolbachia* densities were quantified using single-copy genes of *Wolbachia groEL* and host *rp49*. (A) In abdomens of virgin adult males less than 24 hrs old, CV+ harbor higher *Wolbachia* densities than GF+. (B) In larvae less than 2 days old, *Wolbachia* densities were similar between CV+ and GF+ larvae. *Cytoplasmic incompatibility factor* gene expression for *cifA* and *cifB* was then quantified relatively using *cif* transcripts normalized to *Wolbachia groEL* transcripts. (C-F) *cifA* and *cifB* gene expression in adult males and larvae was similar between CV+ and GF+, indicating *cif* gene expression is uncoupled from CI penentrance variation. (G) Selective cultivation plates for *D. melanogaster* gut bacteria show growth for samples of surface-sterilized fly homogenates of <24-hour-old unmated CV- (white circles) and CV+ males (orange circles) (N=5) in triplicates. For A-F, the p-values are shown for the non-parametric Kolmogorov-Smirnov (and Mann-Whitney U) tests. (H) Gut bacteria CFUs from *Lactobacillaceae* and *Acetobacteraceae* show normalized CFU per male fly from a single plate: *Acetobacteraceae* from MYPL plates and *Lactobacillaceae* from MRS plates. CFUs are markedly higher in CV+ males relative to GF+ males. (I) The CFUs of *Acetobacteraceae* (x-axis) and *Lactobacillaceae* (y-axis) in CV− (white-filled points) and CV+ (orange-filled points) males show a significant positive correlation. Each data point represents the total CFU count per male fly from an individual plate. Regression lines with 95% confidence intervals are shown in grey. A CFU count below 1000 is typical as young flies less than 1 day old have not yet developed a fully established gut microbiome(*50*).

We next investigated if *Wolbachia* reciprocally modify the absolute abundance of the *D. melanogaster* core gut microbiome(*13*). We plated homogenates from CV+ and CV- males onto enrichment and selective medium (Fig 4G)(*49*). *Lactobacillaceae* and *Acetobacteraceae* in CV+ were notably 7.07- and 9.78-fold higher compared to CV-, with loads ranging from approximately 30 to 3,000 CFU per fly (Fig 4H). These values align to previous reports in flies(*50*). Thus, an intact holobiont with both symbiotic classes contains nearly an order of magnitude more cells of the extracellular gut microbiome. Moreover, we observed a significant, positive correlation between the *Lactobacillaceae* and *Acetobacteraceae* abundances (Fig 4I), corroborating prior studies(*51*, *52*). As expected, no CFUs were recovered from negative-control GF+/- (Fig S1). Together, these data indicate that the intact holobiont maintains a substantially higher load of both the gut commensals and *Wolbachia* endosymbiont compared to either symbiotic class alone. Thus, there are likely interorgan, synergistic effects that elevate bacterial densities in the fly holobiont.

### The gut microbiome and endosymbiont independently impact host development and survival

By disentangling the gut microbiome and endosymbiont into constituent parts relative to the intact holobiont, we tested how the gut microbiome and *Wolbachia* differentially impact host fitness (survival) across developmental stages (Fig. 4A-C) and overall embryo-to-adult survival (Fig. 4D). GF+ flies have a modest but significant increase in embryo-to-larvae survival (71.43%) vs. GF- flies (66.95%, Fig 5A). CV+ flies also have slightly, yet significantly, higher embryo-to-larvae survival (87%) than CV- (82%). Thus, *Wolbachia* modestly enhances embryonic survivability. Moreover, CV+ embryos also exhibit significantly higher embryo-to-larval hatching than GF+ (71.43%), indicating that the complete microbiome enhances embryo-to-larvae survivability. From larval-to-pupal and pupal-to-adult stages, *Wolbachia* significantly reduce CV and GF host survival compared to their negative control counterparts, specifying a modest, negative fitness cost at the larval stage onwards (4-13% reduction in survival, Fig 5B,C). Nonetheless, larval-to-pupal survival is significantly higher with a microbiome present (11.67% increase in CV+ vs GF+, 5% increase in CV- vs GF-, Fig 5B), indicating synergism when both classes coexists.

**Fig 5.**
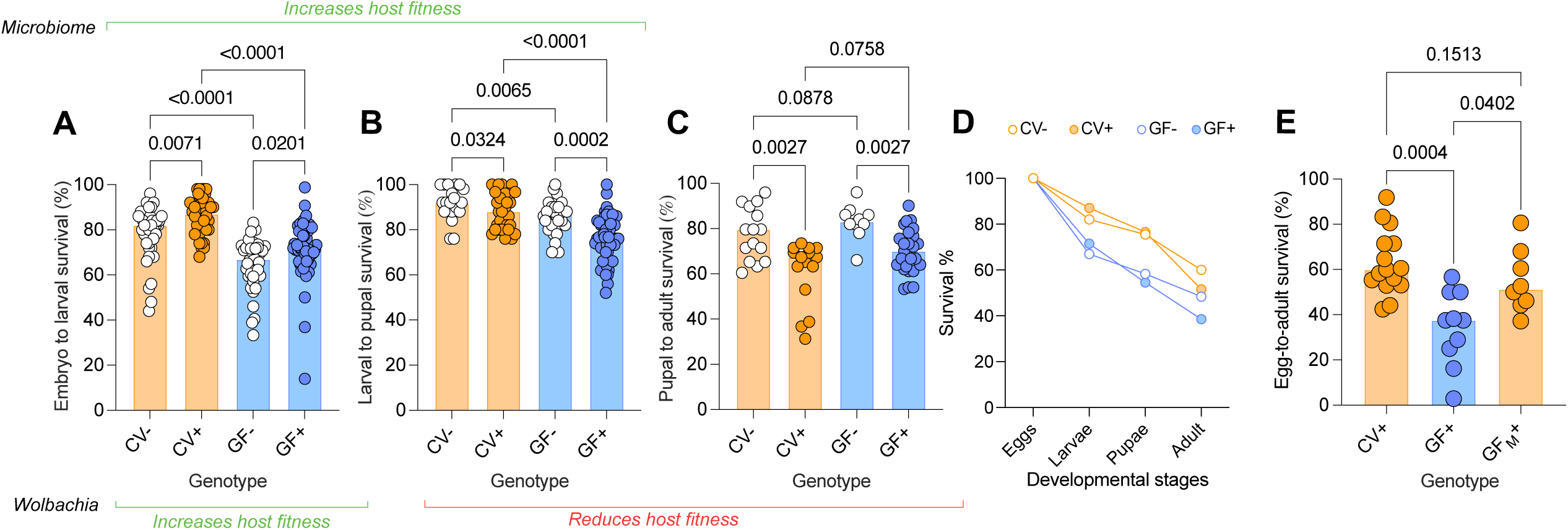
*Wolbachia* and the gut microbiome exert stage-specific, independent contributions to host fitness. To test if the endosymbiont and the microbiome associations affect survival at distinct developmental transitions, we quantified stage-specific survival from (A) embryo-to-larva, (B) larva-to-pupa, (C) and pupa-to-adult. Survival panels reflect the proportion advancing between consecutive stages, whereas (D) the cumulative survival panels show overall developmental outcome. (A) The extracellular microbiome and *Wolbachia* yield independent and positive effects on embryo-to-larval survival, as calcuated by the number of surving larvae divided by the number of emryos laid for CV+ vs. CV- and GF+ vs. GF-. (B) The extracellular microbiome is beneficial to larval-to-pupal survival, while *Wolbachia* imposes a fitness cost in a microbiome-independent manner. (D) Similarly, *Wolbachia* lower pupal-to-adult survival percentage in a microbiome-independent manner. For A-C, the top green bar represents the positive fitness effect of the microbiome on the host, while the green and red lines at the bottom indicate the positive and negative effects, respectively, of *Wolbachia* on the host. (D) To contextualize all survival data, cumulative embryo-to-adult survival plots show GF+ flies consistently performing weker comapred to CV+. (E) When the conventional microbiome is re-introduced into GF+ flies through fecal-to-oral transmission, the egg-to-adult survival of the recipient flies (GF_M_+) significantly increases to the same level as that of CV+ flies. Each circle in the bar charts represents a family, and color denotes genotype status: orange bar (CV), blue bar (GF), open circle (no *Wolbachia*), and filled cirlce (*Wolbachia*). The p-values were calculated with Kruskal-Wallis test followed by Benjamini, Krieger, and Yekutieli multiple correction test.

Overall, GF+/- embryo-to-adult survival (38.56%) was lower than CV+/- survival (51.56%) (Fig 5D) as expected; both values fall within the reported range of 40-75% egg-to-adult survival(*53–56*). Importantly, GF+ egg-to-adult survival (37.5%) was causally attenuated upon a microbiome reintroduction (51.35%) (Fig 5E). Together, these findings specify cumulative impacts of the symbiont classes on survival across development, namely *Wolbachia* are beneficial to embryonic-to-larval development and harmful to subsequent stages, while the gut microbiome increases embryo-to-larval and larval-to-pupal survival during metamorphosis. Such effects may shape host fitness and likely depend on host background(*57*).

## Discussion

Investigations of host-microbe interactions historically oversimplify outcomes as interactions either between hosts and one microbial type or between an anatomical site and its microbial community. However, the impacts of distinct microbial classes in different body sites are neglected. Consequently, holobiont trait variation are commonly attributed to strain, genetic background, and/or environmental variation; and interorgan effects between different microbial classes remain understudied and likely more common than appreciated.

Interorgan effects are often referred to as an “axis” of function. For example, the gut microbiome forms axes with the brain and lung, impacting behavior(*58*) and immune modulation(*59*), respectively. These axes primarily describe the impact of the gut microbiome on other host organs, rather than the functions of the microbial members at those sites. Nascent studies imply the “gut-germline axis” is underappreciated; for example, the gut microbiome impacts murine offspring weight and premature mortality(*60*) as well as worm epigenetic marks in the germline(*61*). These studies overlook multi-partite interactions between different microbial classes across body sites.

Here we address three issues in interorgan, symbiotic interactions that determine offspring survival: (i) Can a reproductive endosymbiont (*Wolbachia*) function autonomously of the gut microbiome? Yes. The endosymbiont’s adaptation is not only self-sufficient from the extracellular microbiome, but stronger in the gut microbiome-free state. This is observed in laboratory fly lines and confirmed in wild-caught flies. (ii) What effects does the gut microbiome impart on densities, penetrance, gene expression, and cell biological underpinnings of the endosymbiont? Mechanistically, CI variation in flies with or without the gut microbiome links to gene expression variation in *Wolbachia* parasitism and disturbed sperm chromatin integrity via increased core histone retention in spermatids and reduced protamine deposition in mature sperm. *Wolbachia* and gut microbial densities also markedly increase in intact holobionts versus hosts lacking one microbial class, implying within-host fitness symbiotic benefits to each other. And (iii) How is host survival shaped by the gut and reproductive microbiome? The two microbial classes cause developmentally-staged and independent impacts on survival. While both classes increase host survival from embryo to larvae; *Wolbachia* modestly reduce viability during subsequent stages of metamorphosis, while the gut microbiome increases survival or has no impact. Importantly, fecal microbiome transplantation causally recapitulates the elevated survival, thus confirming its beneficial role.

If gut-germline effects between microorganisms are widespread, they offer more explanatory depth for understanding phenotypic variation. The challenge is that paired data of microbiomes and functions across body sides are currently insufficient and difficult to collect in practice due to costs and specimen handling needs. Moreover, there is a need for development of holobiont reference databases that couple multi-omic profiles across body sites with bioinformatic approaches for quantitative hologenomics(*3*). Such tools would rapidly speed up identification of the number and types of host and microbial genes and microbial taxa that combinatorially govern phenotypic variation^3^. Indeed, for the CI trait as a case example, CI is well known to vary with *Wolbachia* density, host age, host genetic(*33*, *62–70*). Here, we add a previously unknown dimension of the gut microbiome, and these results are consistent with the gut bacterial genus *Asaia* in mosquitoes can impede vertical transmission of *Wolbachia*(*71*).

In conclusion, this study disentangles the gut microbiome’s impact on a key adaptation. It also unravels the replication and host fitness impacts when the gut and reproductive bacteria are together or alone. The next questions are clear: What core gut microbes influence the reproductive endosymbiont and CI in this system? What are the molecular signals that underpin a gut-germline axis? Is the axis conserved across species, and what is the interplay between this axis and environmental or lifestyle factors? Thus, nested symbiotic interactions between symbionts in different anatomical sites raise a fresh perspective and suite of questions on how one microbial class impacts the functions of another across the biogeography of a holobiont.

## Materials and methods

### Fly genotypes, rearing conditions, and food preparation

*Drosophila melanogaster* y^1^w* (BDSC 1495) lines with *Wolbachia* strain *w*Mel were used for all experiments. *D. melanogaster* white and yellow genetic background is extensively used in cytoplasmic incompatibility (CI) studies(*7*, *33*, *63*, *67*, *72*, *73*). Furthermore, the *Wolbachia* strain present in this line has been effectively utilized in vector control initiatives by the World Mosquito Program(*72*, *74*). *w*Mel uninfected lines were previously established through tetracycline treatment(*33*). *Wolbachia* status was confirmed by PCR using the standard Wolb_F and Wolb_R3 primers (Table S3). Flies were reared in a 12:12 h light/dark cycle at 25°C and 60-70% relative humidity. The food for both CV and GF flies had the same composition and was autoclaved. However, the autoclaved food for GF flies was maintained under sterile conditions, while the food for conventionally reared flies was maintained under non-sterile conditions. The composition for 1L media consisted of 100g yeast, 100g dextrose, 40g cornmeal, 7g agar, and 0.88 mL propionic acid and 7.92 mL of absolute ethanol(*75*).

### Wild *Drosophila melanogaster* collection and isoline establishment

Wild-type *Drosophila melanogaster* were collected from State College, Pennsylvania, USA, using a non-invasive trapping method designed to preserve the natural gut microbiome of field-caught individuals. Traps were constructed by horizontally bisecting a container into two compartments; the lower compartment was filled with a food bait consisting of mashed banana, banana peel, yeast pellets, and approximately 50 mL of water. The bait was covered with a mesh net to prevent direct fly access to the food source. The upper compartment was fitted onto the lower section with lids left open to allow fly entry. Flies were monitored daily throughout the collection period. This trapping design ensured that the natural fly microbiome remained unperturbed, as flies were neither exposed to the bait substrate nor subjected to prolonged starvation. Following collection, flies were transported to the laboratory in germ-free food to prevent microbial contamination from the laboratory food during transit. Upon arrival, females were immediately inspected for mite infestation. Each female was then individually housed in a germ-free vial to establish isogenic lines. After approximately one week, when F1 pupae had formed, the founding female was used to assess *Wolbachia* infection status and confirmation of *Drosophila melanogaster* species using COX1 sanger sequencing. Flies were subsequently reared for an additional generation to obtain sufficient numbers for egg collection, conventional/germ-free treatment, and hatch rate assays.

### Rearing of flies to obtain mothers and fathers for hatch rate assay

Adult flies of both sexes were kept for oviposition and cleared on the 5^th^ day. The virgin female offspring (referred to as PGMs – paternal grandmothers) were collected on the 11^th^ to 13^th^ day. The PGM virgins were kept in isolation for 4-5 days to ensure virginity and to obtain moderate CI penetrance to test if germ-free treatment yields higher or lower CI penetrance(*62*). Virgin PGMs were crossed with males to obtain embryos. In brief, ∼30 virgin PGMs, aged 4-7 days, were allowed to mate overnight with 10-30 males in an empty bottle secured with a grape plate smeared with yeast. The next morning, the grape plate was replaced by a fresh one and incubated for 30 min. The grape plate was again replaced by a new one and incubated for ∼4 hours to obtain sufficient embryos. In all instances, the same amount of yeast was smeared with a spreader on the grape plate to stimulate egg laying. All eggs from each grape plate within a treatment are collected in a petri dish with milli-Q water to ensure homogeneity. This also diluted the yeast, and thus the egg collection technique does not introduce extra yeast into the system. The embryos were processed for conventional (CV) and germ-free (GF) treatments^51^. The GF treatment was carried out as previously described(*76*). Briefly, embryos were dechorionated twice using 10% bleach, followed by a single wash in 70% ethanol and three washes in PBS containing 0.1% Triton X (Fig S4). On the 2^nd^ and 9^th^ days after the CV and GF treatments, larvae and pupae survival were counted, respectively. On the 10^th^ to 13^th^ day, adult emergence was noticed. Males (referred to as fathers) less than 24 hours old were collected to perform hatch rate assay. Meanwhile, parental flies of both sexes were reared on food for 4 days to lay fertilized eggs, after which they were removed. The F1 virgin female offspring (referred to as mothers) were collected on the 10^th^ to 13^th^ day.

### CI assays

All crosses were performed using <1-day-old males because they cause strong CI lethality(*77*) and 3-5-day-old females in a conventional incubator under a 12:12 h light/dark cycle at 25°C with 60–70% relative humidity^16^. The matings were performed in an 8 oz polypropylene bottle and secured with a grape plate. The flies were kept overnight, and the grape plate was replaced by a fresh new grape plate. After 24 hours, the flies were cleared, and the number of embryos on the grape plate was counted (referred to as the 0-hour count). The embryos were incubated for an additional 27 hours, and the unhatched embryos were counted (referred to as 27-hour count). Any plate with less than 25 embryos was discarded. The hatching percentage was determined by calculating the percentage of embryos hatched on the 27-hour count compared to the 0-hour count. CI (incompatible) crosses were performed between *Wolbachia*-infected males (CV+ and GF+) and *Wolbachia*-uninfected females (CV-). Compatible rescue crosses were performed between *Wolbachia*-infected males (CV+ and GF+) and *Wolbachia*-infected females (CV+). Using CV+ females for rescue experiments serves as a persistent control that ensures consistency in rescue studies(*7*, *28*, *33*, *62*, *65*, *67*) and avoids confounders by adding the extra variable of GF+ females.

### Germ-free check and CFU quantification

To score the bacterial CFU in CV flies or to determine the germ-free status in GF flies, less than 24 hours old adult male flies (N=5) from each treatment were collected. The flies were surface sterilized by washing twice in 70% Ethanol and once in 1x Phosphate-buffered saline (PBS). The surface sterilized flies were crushed with a sterile micro pestle, diluted in 300 uL 1xPBS, and 20 uL was used to spread onto MYPL (*Acetobacteraceae* selective medium)(*49*) and MRS agar (*Lactobacillaceae* enrichment medium) (Sigma, #69964, USA). The MYPL media was composed of 1% D-mannitol (Sigma, #M4125, USA), 1% yeast extract (VWR, #J850, USA), 0.5% peptone (Thermo Fisher Scientific, # R451102, USA), 1% lactic acid (Sigma, #L6661, USA), 0.1% tween 80 (Thermo Fisher Scientific, #278632500, USA), adjusted to pH 7 with NaOH pellets (Spectrum Chemical, #S1301, USA), and 1.5% agar (Sigma, #A1296, USA) was added. The plates were incubated for 48 hours at 30°C and the CFU per fly was quantified using the equation:

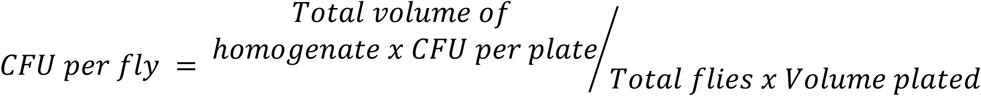

### Microbiome reintroduction

A natural method of gut microbiome transfer based on fecal material ingestion followed a previously published protocol(*78*). First, 50 adult *w*Mel+ males of random age were left in a GF vial for 7 min. Second, vials were flipped upside down for another 7 min so that the fecal material was deposited onto the plug and food surface. Finally, the resulting vials were externally surface sterilized by spraying them with ample 70% EtOH before adding the GF embryos to grow and ingest the bacteria.

### Gnotobiotic Fly Establishment

To cultivate and use the core commensal microbiota of the flies, five males and five females were surface-sterilized and homogenates were plated onto MRS and MM agar plates (see Germ-Free Check and CFU Quantification). Unique colonies were enriched in their respective liquid media and identified by Sanger sequencing of the 16S rRNA gene. The identified members of *Acetobacteraceae* and *Lactobacillaceae* were subsequently used to prepare three bacterial fractions: live cell pellet, cell-free supernatant, and heat-killed pellet. For the heat-killed pellet, fresh overnight cultures were adjusted to 5×10⁵ CFU/50 µL, heat-killed at 60°C for 45 minutes using a heat block(*79*), centrifuged at 10,000 rpm for 3 minutes, and the resulting pellet was resuspended in 1 mL of fresh sterile liquid media. For the cell-free supernatant (CFS) preparation, fresh cultures adjusted to 5×10⁵ CFU/50 µL were centrifuged at 10,000 rpm for 3 minutes, and the supernatant was collected and subjected to a second centrifugation step to remove residual cells, followed by 0.2 µm syringe filtration to eliminate large debris or live cells while retaining metabolites and perhaps viruses, and readjusted to a final volume of 1 mL using sterile liquid media; the retained pellet from this step was resuspended to 1 mL in sterile media to serve as the live cell pellet treatment. For all three preparations, 50 µL was used to inoculate onto germ-free eggs per vial, and 20 µL of each preparation was plated onto MM and MRS agar plates to confirm bacterial viability or inviability as appropriate.

### RNA/DNA isolation for RT/qPCR and transcriptome

RNA was extracted and used for RT-qPCR (N=6 to 12 replicates) and RNA-Seq (N=3 replicates) from samples collected at various developmental stages for each treatment: 2-day-old larvae (N=7), late-stage pupae (N=1), and less than 24-hour-old adults (N=1). Kl-5 primers were used to sex individual pupae collected at the late stage(*80*). PCR conditions used were 95°C for 2 min and 30 cycles of 95°C for 30 sec, 56°C for 30 sec, and 72°C for 1 min, final extension of 72°C for 5 min. Whole body larvae/pupae/adults were frozen in liquid nitrogen and crushed with a sterile micro pestle. For *Wolbachia* density measurements, the adult abdomen was dissected and processed. RNA was extracted using 1 mL TRI Reagent (Sigma, #93289, USA) and 200 uL chloroform (Sigma, #C2432, USA). Samples were incubated for 3 min in an ice block with occasional mixing and centrifuged for 15min/12000rpm/4°C. The upper aqueous phase was transferred to an RNase-free microcentrifuge tube containing 500 uL isopropanol & 1 uL glycogen. Samples were incubated for 10 min in an ice block with occasional mixing, and centrifuged for 10min/14000rpm/4°C. The supernatant was discarded, and the pellet was washed in 500 uL 75% EtOH followed by centrifugation for 5min/7500rpm/4°C. The pellet was air-dried inside the laminar flow hood at room temperature (RT) until it became translucent. The RNA pellet was resuspended in 30 uL of nuclease-free water and incubated at 55°C for 15 min for dissolution. Any contaminating DNA was eliminated by treating with DNA-free™ DNA Removal Kit (Thermofisher Scientific, #AM9937, USA) as per the manufacturer’s instructions. RNA was confirmed DNA-free through negative PCR amplification of *rp49 Drosophila* gene. DNA-free RNA samples were then purified using the ethanol precipitation method by initially raising to 100 uL using nuclease-free water. 10 uL of 3M Sodium Acetate (Thermofisher Scientific, # R1181, USA) and 300 uL of 100% EtOH were added in the sample, mixed thoroughly & incubated at - 80°C overnight. The next morning, samples were centrifuged at full speed (14000 rpm) for 30 min at 4°C and supernatant was discarded. 500 uL of freshly prepared 70% EtOH was added and centrifuged at full speed for 20 min at 4°C. The supernatant was discarded and pellet was air dried for 30 min inside the biosafety cabinet until it became transparent. Pellet was resuspended in 20 uL nuclease-free water and incubate at 55°C for 15 min to fully dissolve. The purified RNA sample was stored at -80°C until further use. cDNA was synthesized from RNA (100 ng for larvae and 1 ug for adults) with SuperScript VILO kit (Invitrogen, #11755250) and diluted 1:10 using Ambion nuclease-free water.

DNA was isolated from the non-aqueous layers from the samples used for RNA isolation using Qiagen PureGene Core Kit A kit (Qiagen, #158667, USA) following manufacturer’s instructions. However, during the DNA precipitation step we added 500 uL of 100% Isopropanol and 1 uL of glycogen to enhance the DNA recovery. qPCR was performed in 2 technical replicates using 1 uL of the diluted cDNA or DNA (10ng) and PowerTrack SYBR Green Mastermix (Applied Biosystems, #A46109) on a QuantStudio 6 Pro Real-Time PCR system using the following conditions: 50°C 2 min, 95°C 10 min, 40 cycles (15 s at 95°C, and 30 s at 56°C and 60°C for whole body and abdomen samples, respectively), 95°C 15 s, 60°C 1 min, 95°C 15 s. Gene expression of *cifA* and *cifB* (Table S3) was normalized to *groEL* and *Wolbachia* density was measured by 2^-ΔCt^ method using *groEL* relative to *rp49*. Ct values above 30 were omitted from the analysis.

### Transcriptome

Samples were prepared by generating a uniquely indexed library from each of the RNA samples isolated from male pupae using the Illumina Stranded Total RNA Prep with RiboZero Plus Microbiome kit (Illumina, USA) and custom *D. melanogaster* probes for the rRNA depletion. The libraries were pooled into a single pool containing an approximately equimolar amount of the samples and outsourced to the Genomic Sciences Laboratory, North Carolina State University, for sequencing using NovaSeq6000 S4 150 PE lane yielding ∼25 million pairs of reads per sample on average. The raw reads were subjected to quality control using Trimgalore v0.6.10 with a PHRED score of 20 and minimum read length of 25 bp. The cleaned reads were mapped to *D. melanogaster* reference genome (v.6.12) using STAR v2.7 to generate BAM files. The read counting was performed using the BAM files with featureCounts v2.0.6. The raw reads were rarefied using EcolUtils v0.1 with 100 permutations setting the lowest read (16M) as the threshold. The reads were then processed for downstream differentially expressed genes (DEGs) using DeSeq2 v1.42.0. Data deposition: The data reported in this paper have been deposited in Sequence Read Archive (SRA; https://www.ncbi.nlm.nih.gov/sra) (accession no. PRJNA1209084).

### Histone staining and quantification

Less than 8 hours old males from the same population used for hatch rate assay above were dissected to collect testes in chilled 1x PBS solution and perform histone staining as described previously(*7*). In brief, the testes were fixed for 30 min at RT in 4% formaldehyde prepared in 1x PBS with subsequent washing with 1x PBST solution (1x PBS + 0.3% Triton X-100) and blocking in 1% BSA for core histone and 3% BSA for H4ac (h4 acetylation) in PBST at RT for an hour. Samples were incubated overnight with core histone antibody (MilliporeSigma, Cat#MABE71, USA) (1:1,000 dilution in BSA-PBST) at 4°C in rotating shaker. After washing with 1x PBST solution (1x PBS containing 0.1% Triton X-100), samples were incubated with goat anti-mouse Alexa Fluor 488 secondary antibody (Thermo Fisher Scientific, Cat#A28175, 1:500 diluted in 1x PBST) for core histone or Goat anti-rabbit Alexa Fluor 594 secondary antibody (1:500 diluted in 1x PBST) for H4ac in the complete dark at RT for 4 hours. After washing in 1x PBST, samples were then mounted in vectashield DAPI-containing mounting media (Vector Laboratories, Cat#H-1200-10, USA) for imaging using Zeiss LSM 880 confocal microscope with constant exposure settings across treatment groups. The signal intensities from early canoe stage sperm bundles that retained the H4ac signal and late canoe-stage sperm bundles that retained core histones signal were quantified as previously described(*7*), and plotted in GraphPad Prism.

### Protamine staining and quantification

Protamine deficiency was evaluated using chromomycin 3 (CMA3) stain as described previously(*7*). In brief, mature sperm were extracted from seminal vesicles of 1-2-day-old virgin males in chilled 1x PBS. Samples were fixed in 3:1 vol/vol methanol:acetic acid for 20 min at 4°C. Subsequently, the slide was air-dried and treated with 0.25 mg/mL of CMA3 in McIlvain’s buffer, pH 7.0, with 10 mM MgCl_2_ for 20 min. The samples were washed with 1x PBS and mounted in vectashield DAPI-containing mounting media. Imaging was performed using Zeiss LSM 880 confocal microscope. Fluorescence quantification was performed by scoring fluorescent pixels in arbitrary units (A.U.) within individual sperm heads using ImageJ. The flies were reared at 21°C instead of 25°C for this assay.

### Statistical analysis

Comparisons between two groups were performed using Mann–Whitney U tests (Wilcoxon rank-sum tests). Comparisons among three or more groups were performed using the Kruskal–Wallis test, followed by Dunn’s post-hoc test for pairwise comparisons when the Kruskal–Wallis test was significant. P values were adjusted for multiple comparisons using the false discovery rate (FDR) method. Outliers in the hatch rate assay evaluating *eci* gene overexpression (WD0633) were identified using the 1.5 × interquartile range (IQR) criterion, with values below Q1 − 1.5 × IQR or above Q3 + 1.5 × IQR excluded from the corresponding analyses. Data were visualized using the ggplot2 package (version 4.0.3).

## Acknowledgments

The authors thank the Huck Institutes Genomics Core Facility (RRID:SCR_023645) at The Pennsylvania State University for generating a uniquely indexed library from each of the RNA samples and custom *D. melanogaster* probes for the rRNA depletion. The co-authors also thank the Microscopy Core Facility (RRID:SCR_024457) for use of the Zeiss LSM 880 confocal microscope. The authors also thank Will Ludington’s lab, Department of Embryology, Carnegie Institution for Science, Baltimore, MD, USA, for their expert guidance in generating germ-free flies and proofreading of the manuscript. Research reported in this publication was supported by NIH R01 AI179743, NIH R01 AI189624, and Pennsylvania State University to SRB. Approximately 60% of the total project costs was financed with Federal funds from two NIH awards totalling $1,946,715 and $1,153,655 to date. The remainder was funded by university fund sources. The content is solely the responsibility of the authors and does not necessarily represent the official views of the National Institutes of Health.

## Author contributions

M.I. and S.R.B conceived the project. M.I. and S.R.B., Sa.R.B., R.K., and S.L.F. designed the research. M.I., S.L.F, R.K, D.H.Z.L, M.W.B, E.L, A.M., L.R.C., and C.L. performed the research. M.I., R.K, E.L, S.R.B., and Sa.R.B. analyzed the data. M.I. and S.R.B wrote the manuscript. M.I. and S.R.B. revised the manuscript.

## SUPPORTING INFORMATION

**S1 Fig.**
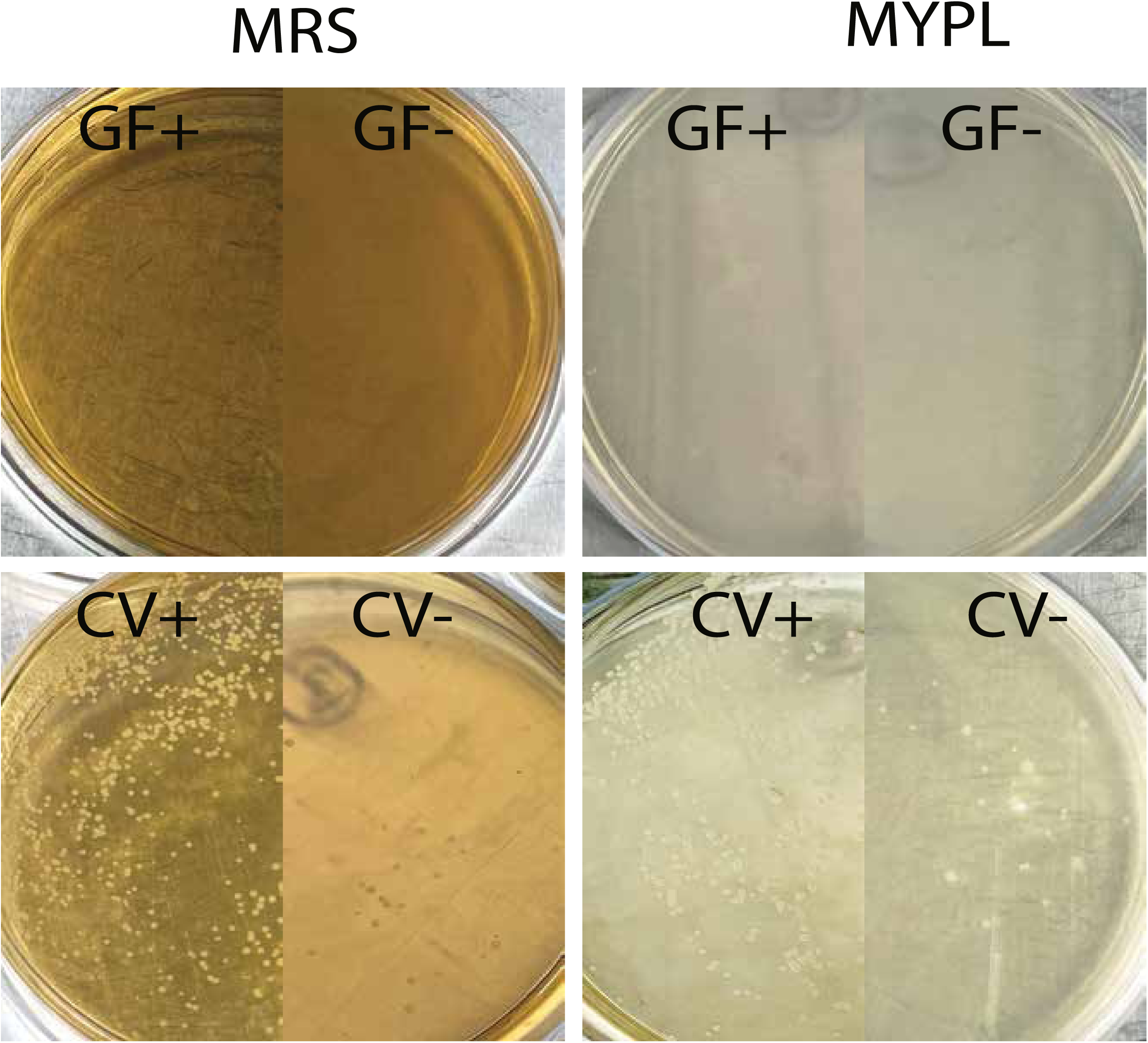
Confirmation of gut microbiome absence and presence by colony forming units (CFUs) in GF and CV lines. The germ-free status of the GF-/+ lines was confirmed by the absence of colonies when 20uL of fly homogenate were plated on to MYPL and MRS plates. The conventionally-reared positive control lines had ample colonies with colony number significantly elevated in CV+ relative to CV- flies.

**S2 Fig.**
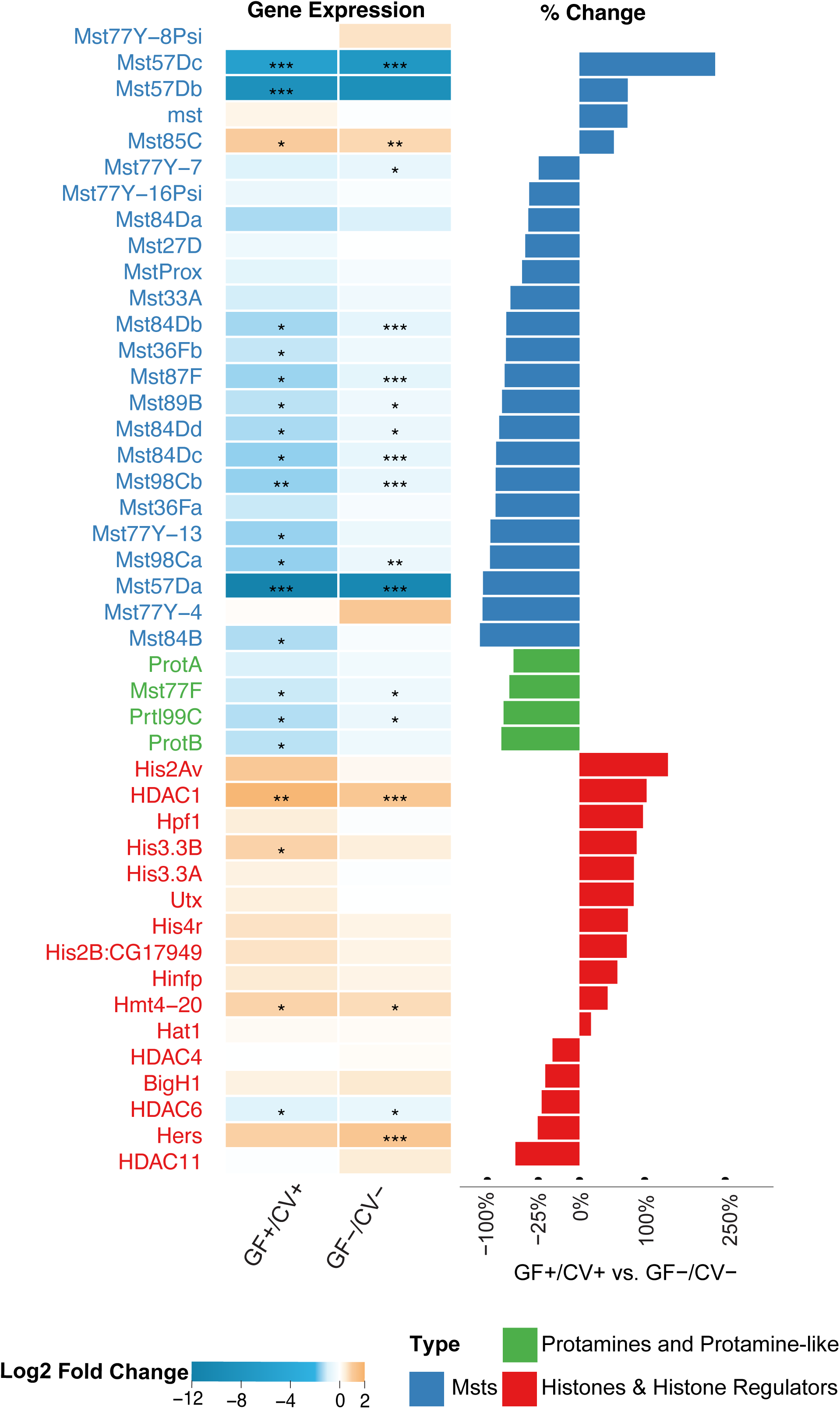
Host regulation of histone, protamine, and male-specific transcripts. Comparative transcriptomic analysis of GF+ vs. CV+ male pupae and GF- vs. CV- male pupae revealed significant differentially expressed genes (Table S2) narrowed here to 38 genes expressing histone, protamine, and male-specific transcripts (MSTs). Notably, many histone-related genes genes (red) were upregulated (consistent with CI-defining histone retention), while protamine (green) and MST (blue) genes were downregulated (consistent with CI-defining protmaine deficiency). The bar chart shows the percent change difference between GF+/CV+ versus GF-/CV-. Log2 fold change was calculated using DESeq2, and its default parameters; Walt test and Benjamini-Hochberg procedure were used to calculate p and adjusted p-values.

**S3 Fig.**
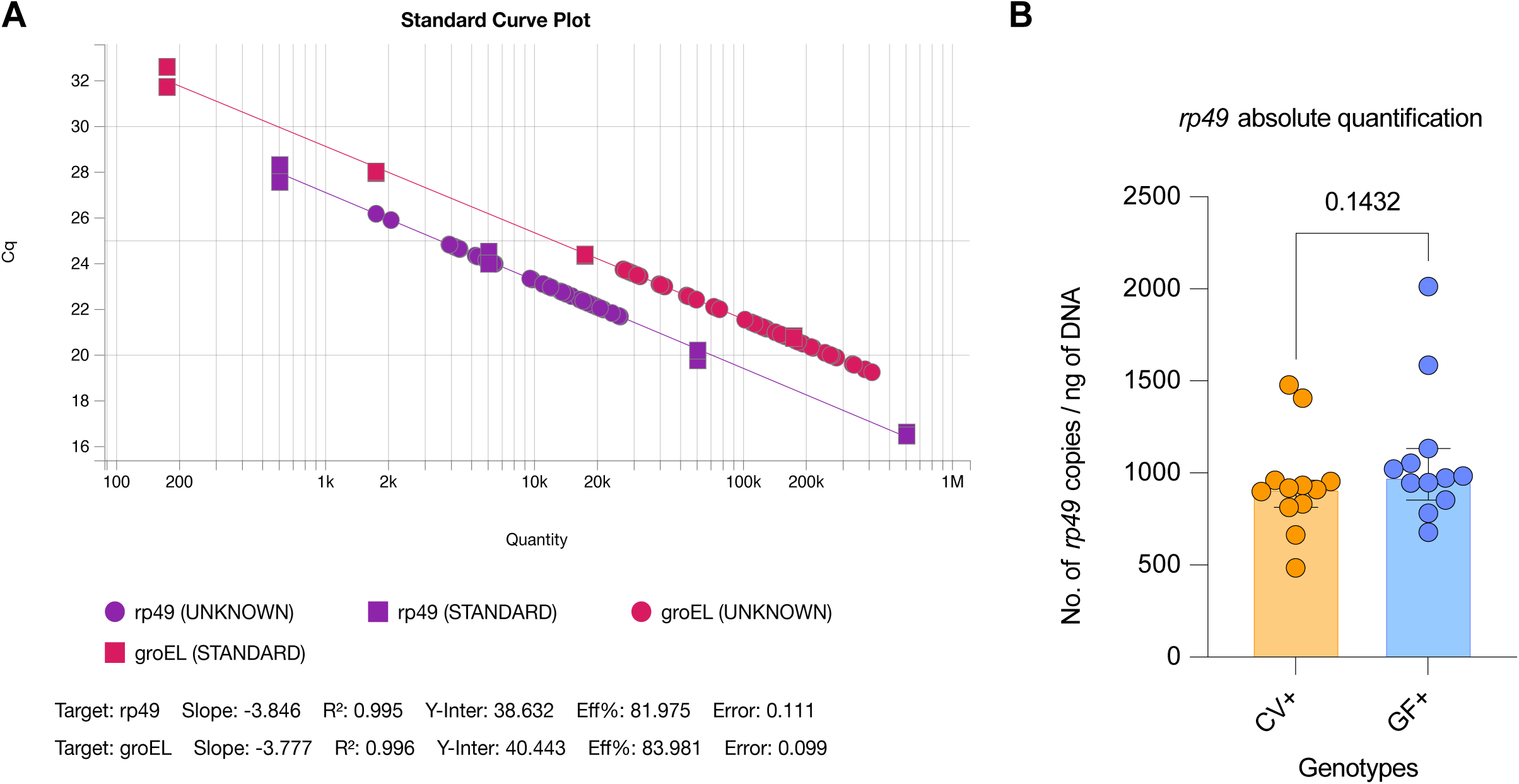
Copy number of host *rp49* in CV+ and GF+ are comparable. (A) Standard curves for the absolute quantification of *rp49* were generated using a log10 dilution series of known amounts of PCR product amplified from a larger region of the same *rp49* gene (404 bp). PCR amplification was performed using primers listed in Table S1 at a concentration of 0.4 µM. The cycling conditions were as follows: 30 cycles of 95°C for 15 seconds, 60°C for 30 seconds, and 72°C for 30 seconds. The PCR products were purified using the Monarch DNA Purification Kit (NEB) and their concentrations were measured using a NanoDrop One spectrophotometer. The number of *rp49* copies was calculated using the following formula: Number of DNA copies = (amount of DNA in ng * 6.022 x 10^23) / length of amplicon in bp x 10^9 x 660). Standard curve was plotting using Design & Analysis Software v2.8.0 (B) Mann Whitney U test of the CV+ and GF+ shows that the absolute count of CV+ and GF+ *rp49* gene in their abdomen are similar.

**S4 Fig.**
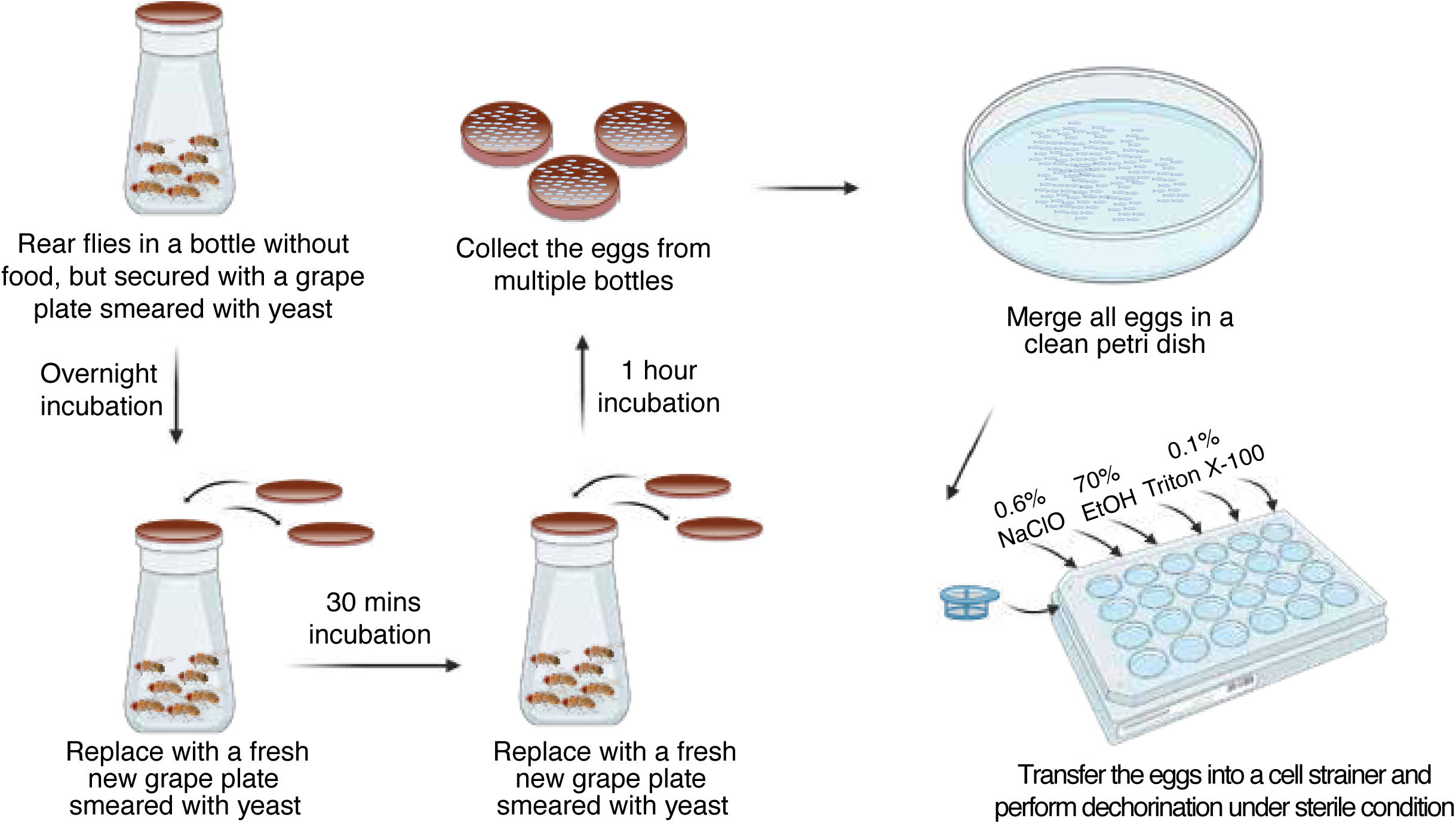
Dechorination of *Drosophila melanogaster* embryos. The germ-free treatment followed the standard procedure of bleaching fertilized embryos. Briefly, CV- and CV+ females were allowed to oviposit overnight on a grape plate smeared with yeast. The next morning, the grape plates were replaced with fresh ones smeared with yeast, and the flies were allowed to oviposit for 30 minutes. This was followed by another replacement with a fresh grape plate smeared with yeast, where the flies oviposited for an additional hour to collect 1-hour embryos. The collected embryos were transferred to a petri dish and approximately half was transferred into a cell strainer. These embryos were bleached twice in 10% bleach for 2.5 minutes each, followed by a single wash in 70% ethanol for 30 seconds and three quick washes in PBS containing 0.1% Triton X for 10 seconds each. The dechorionated eggs were carefully counted under a stereomicroscope and then placed on a germ-free grape plate or a germ-free food vial before being incubated under sterile conditions.

**S1 Table:**
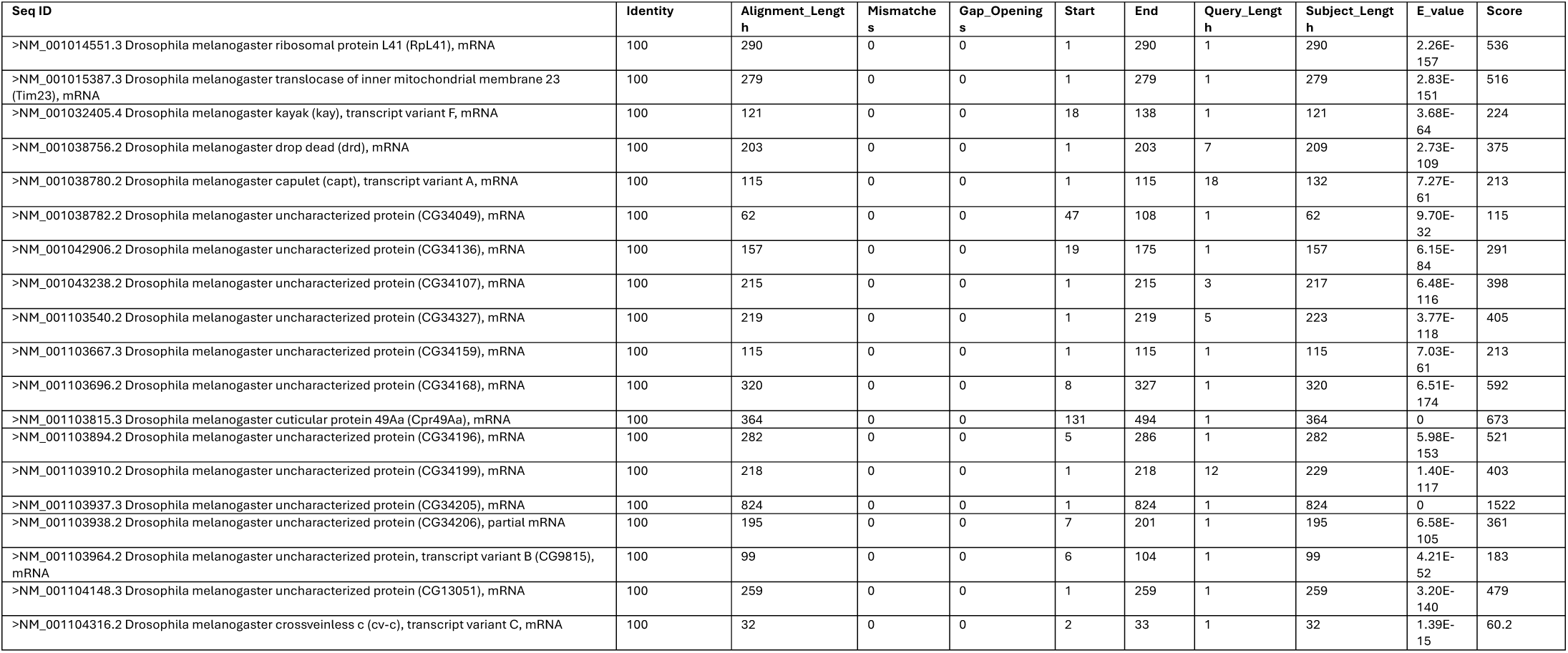
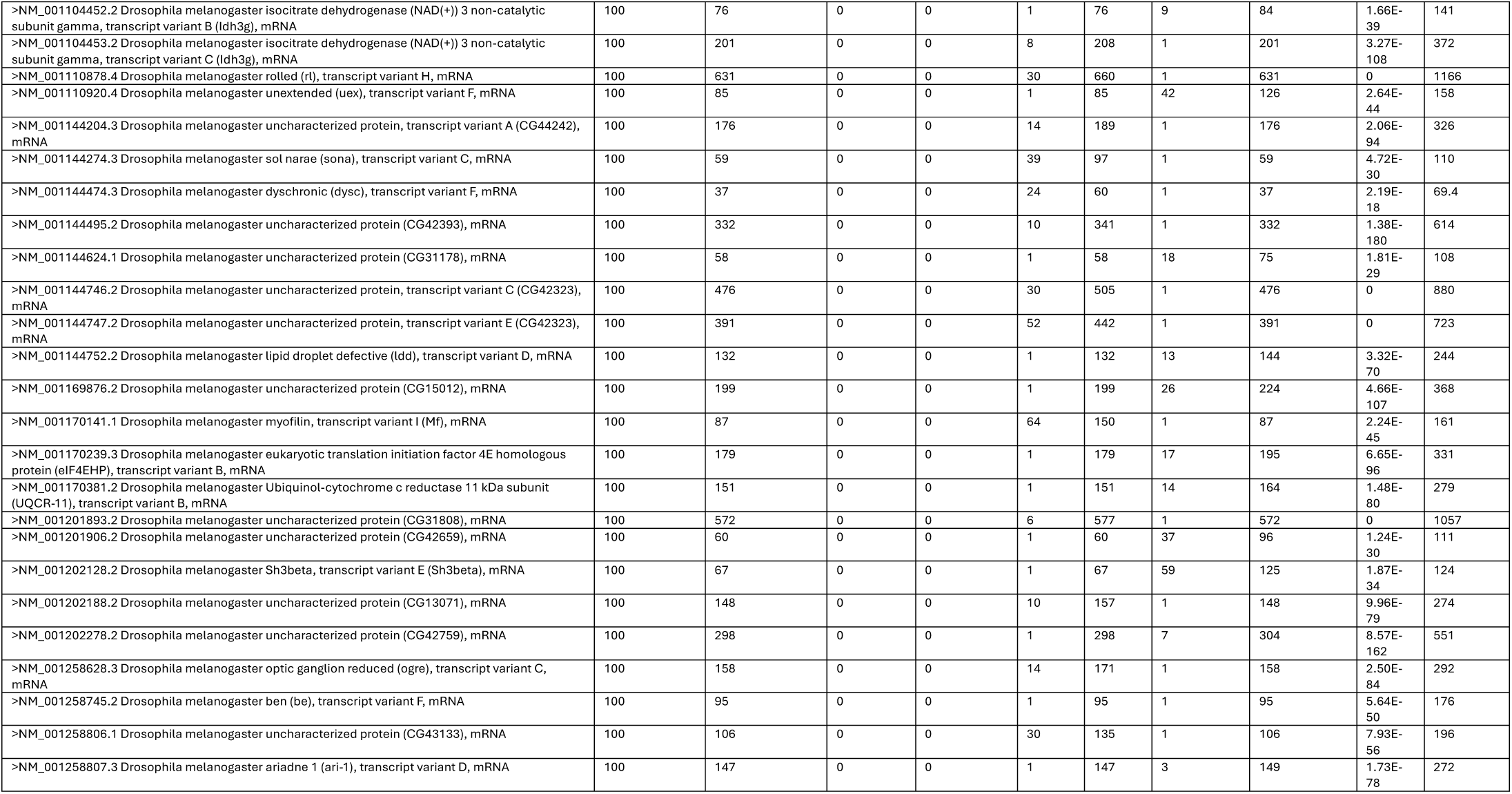
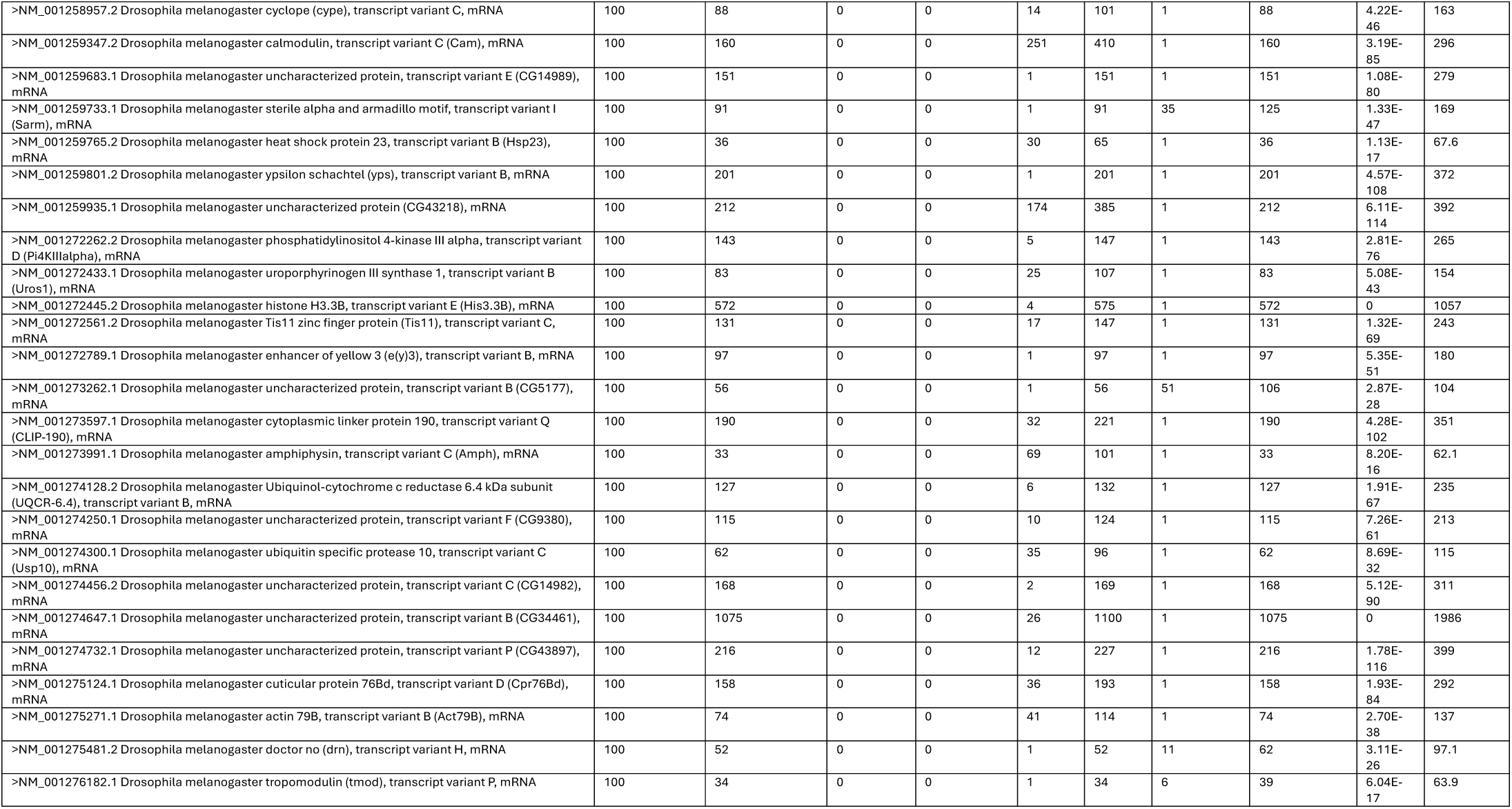
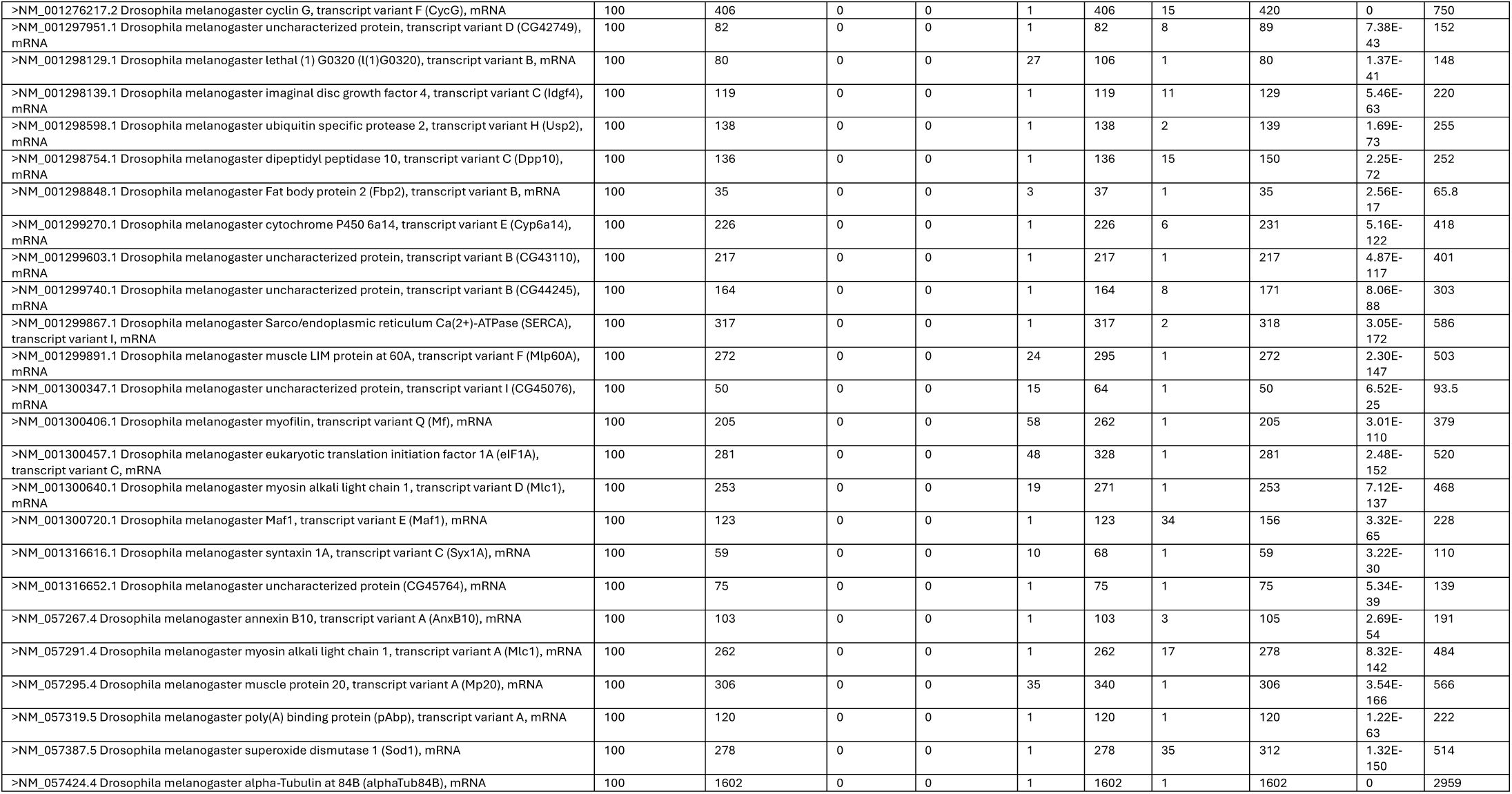
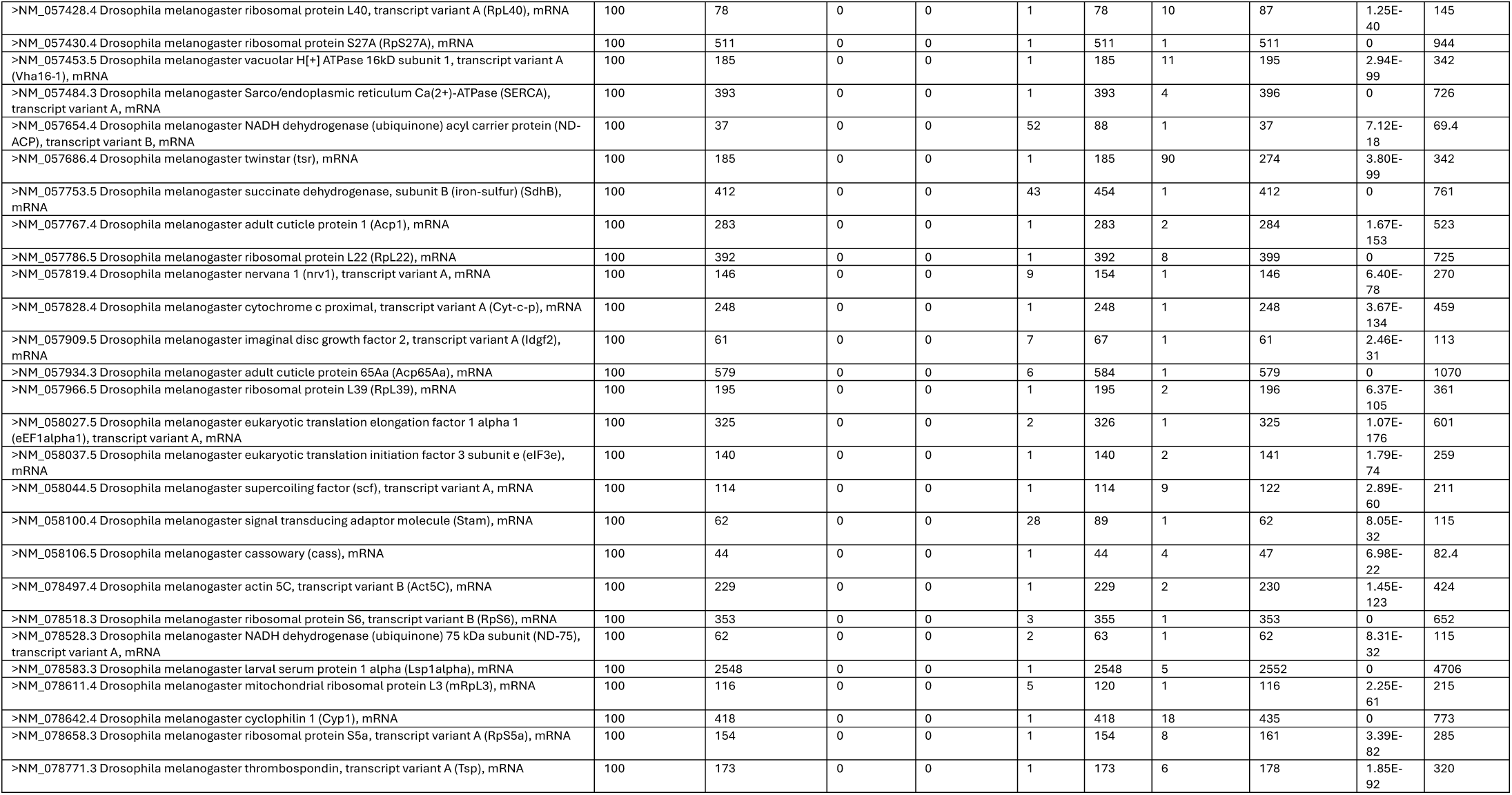
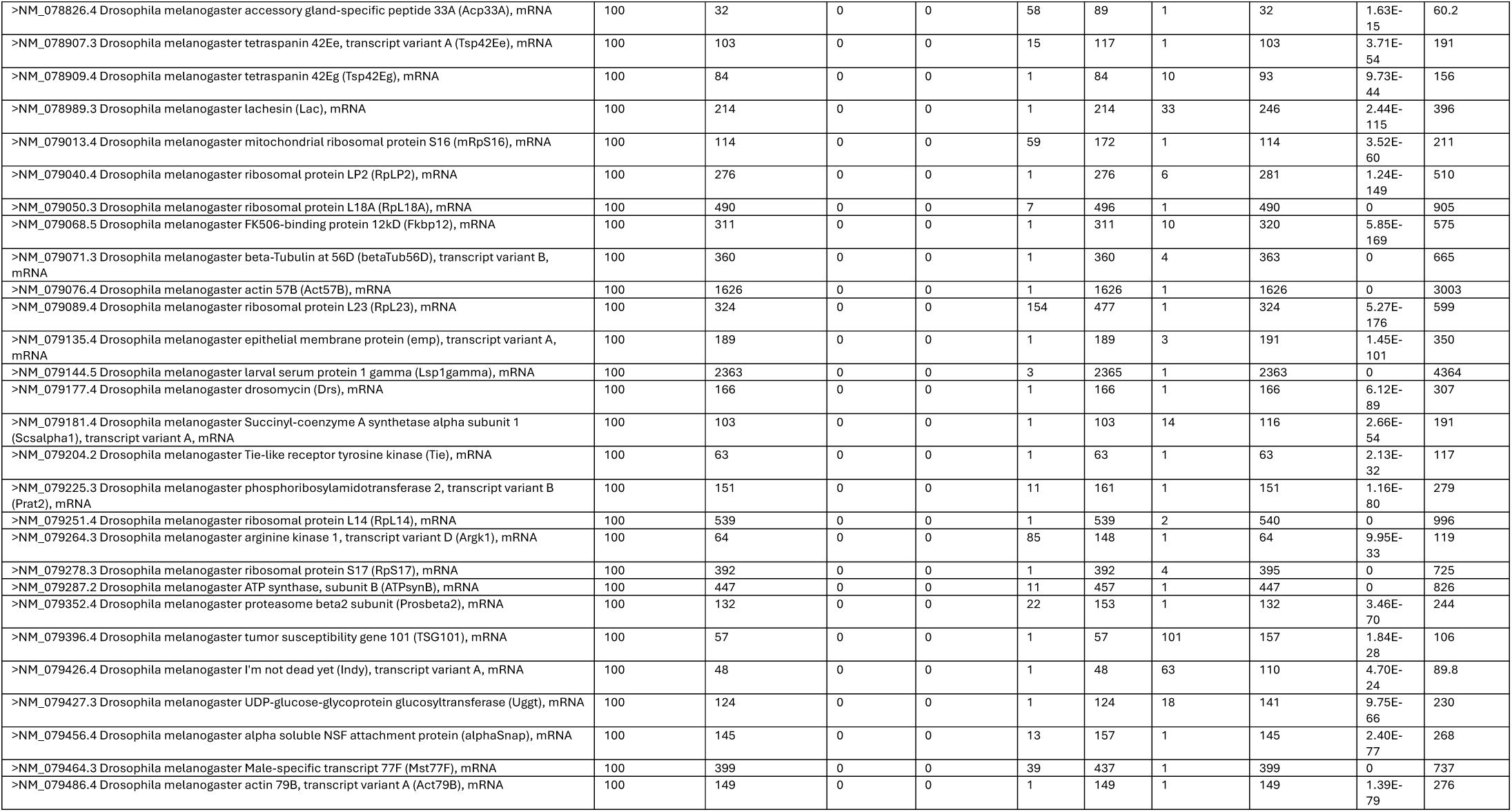
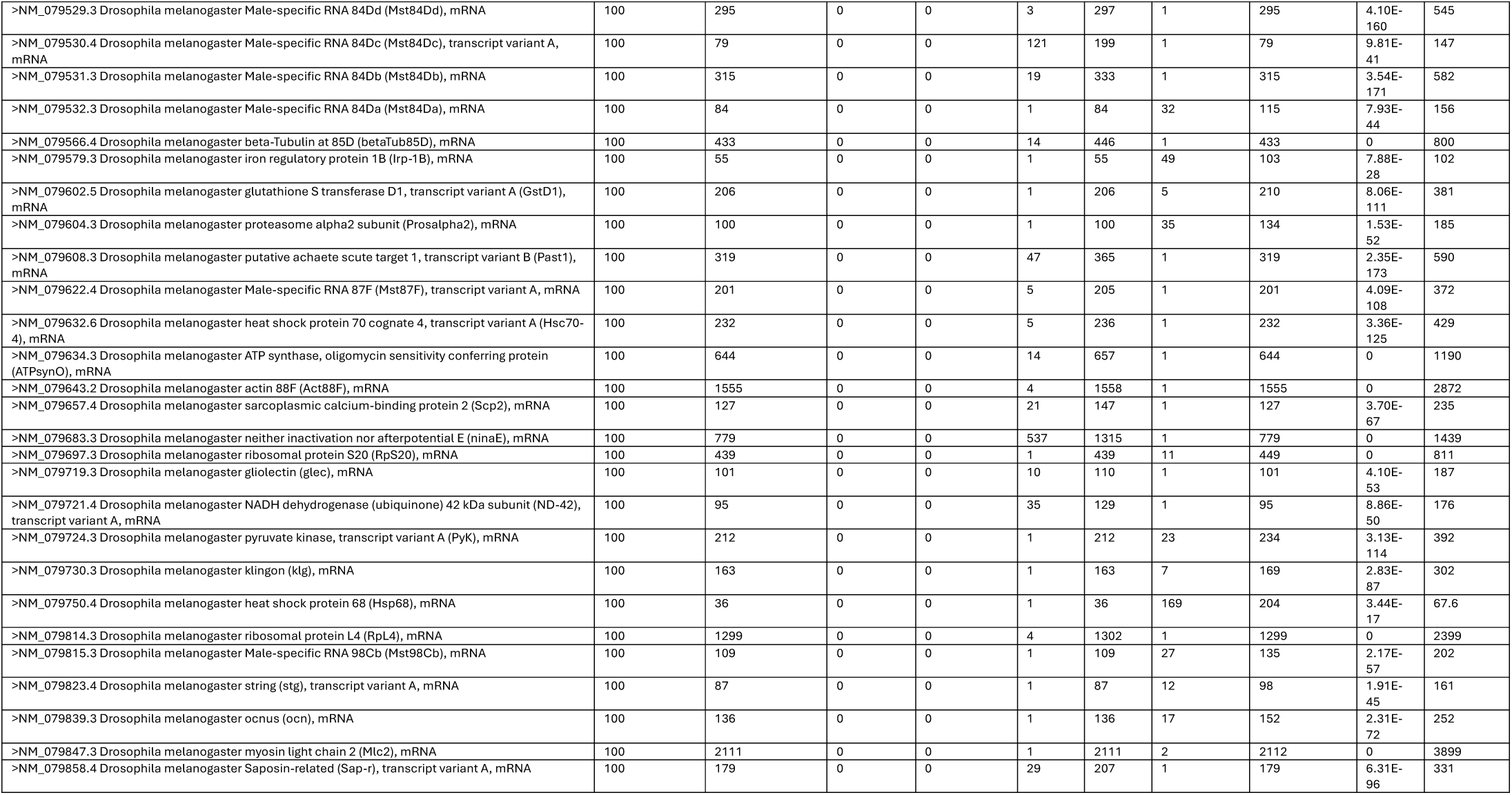
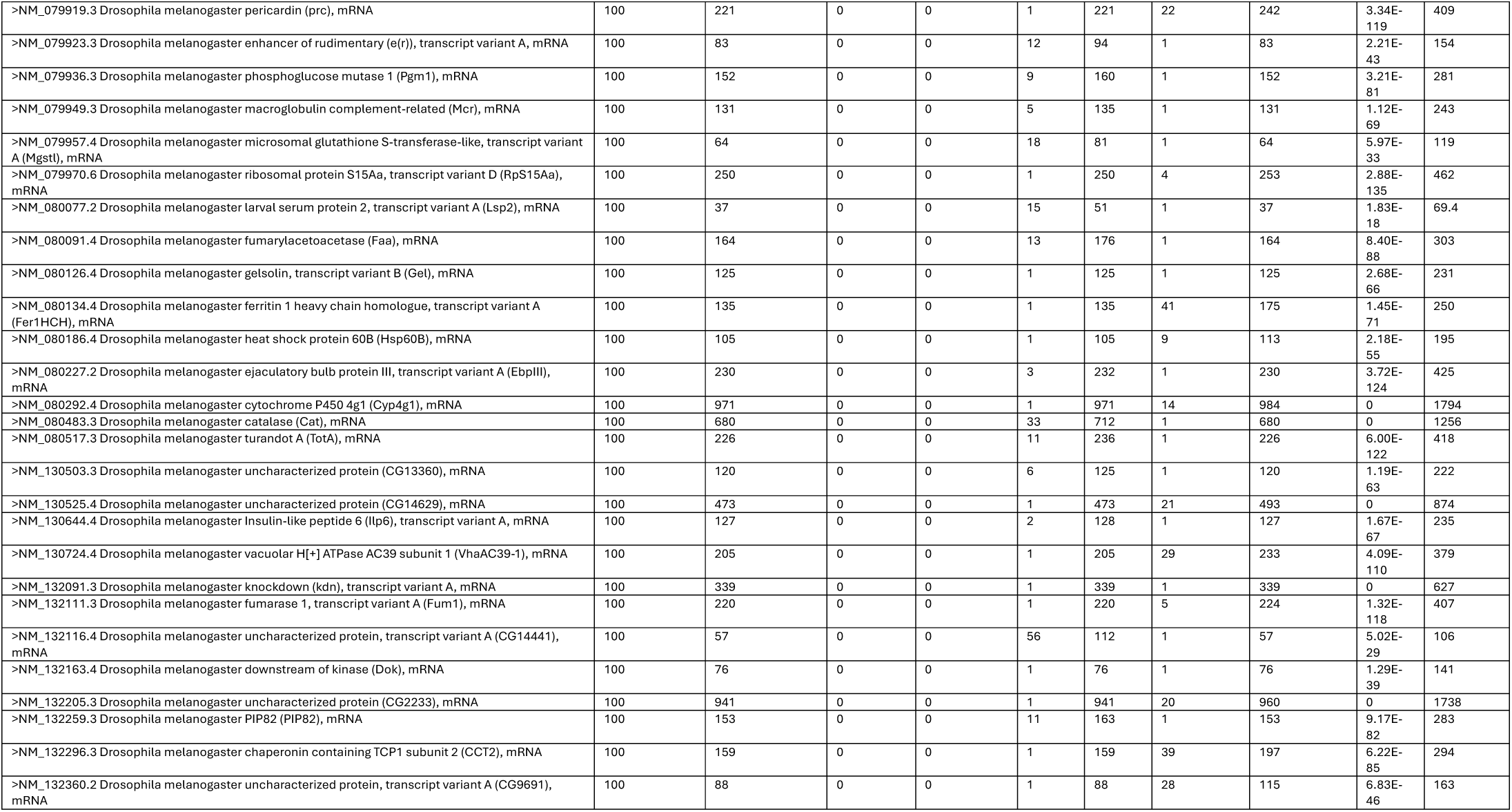
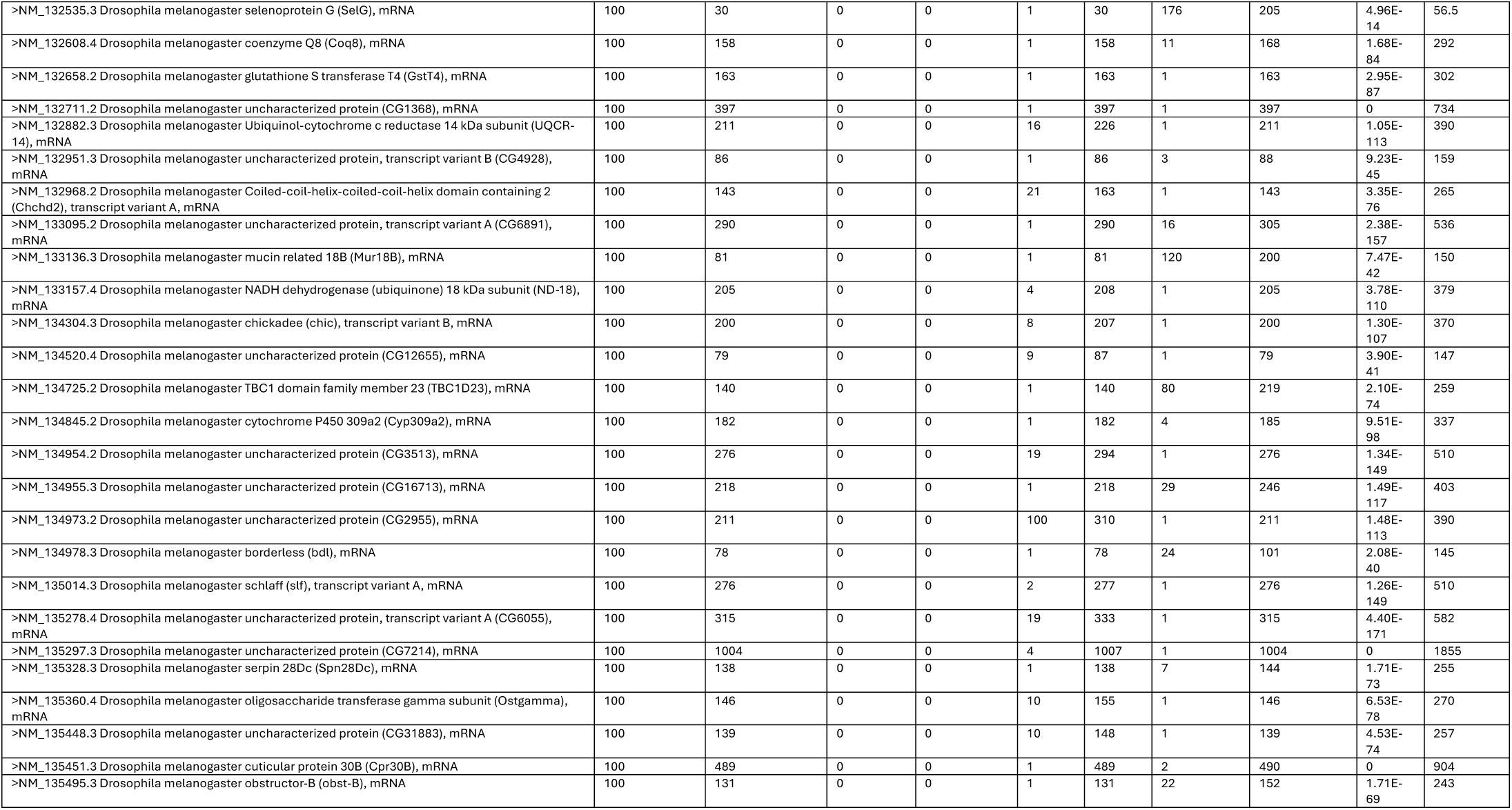
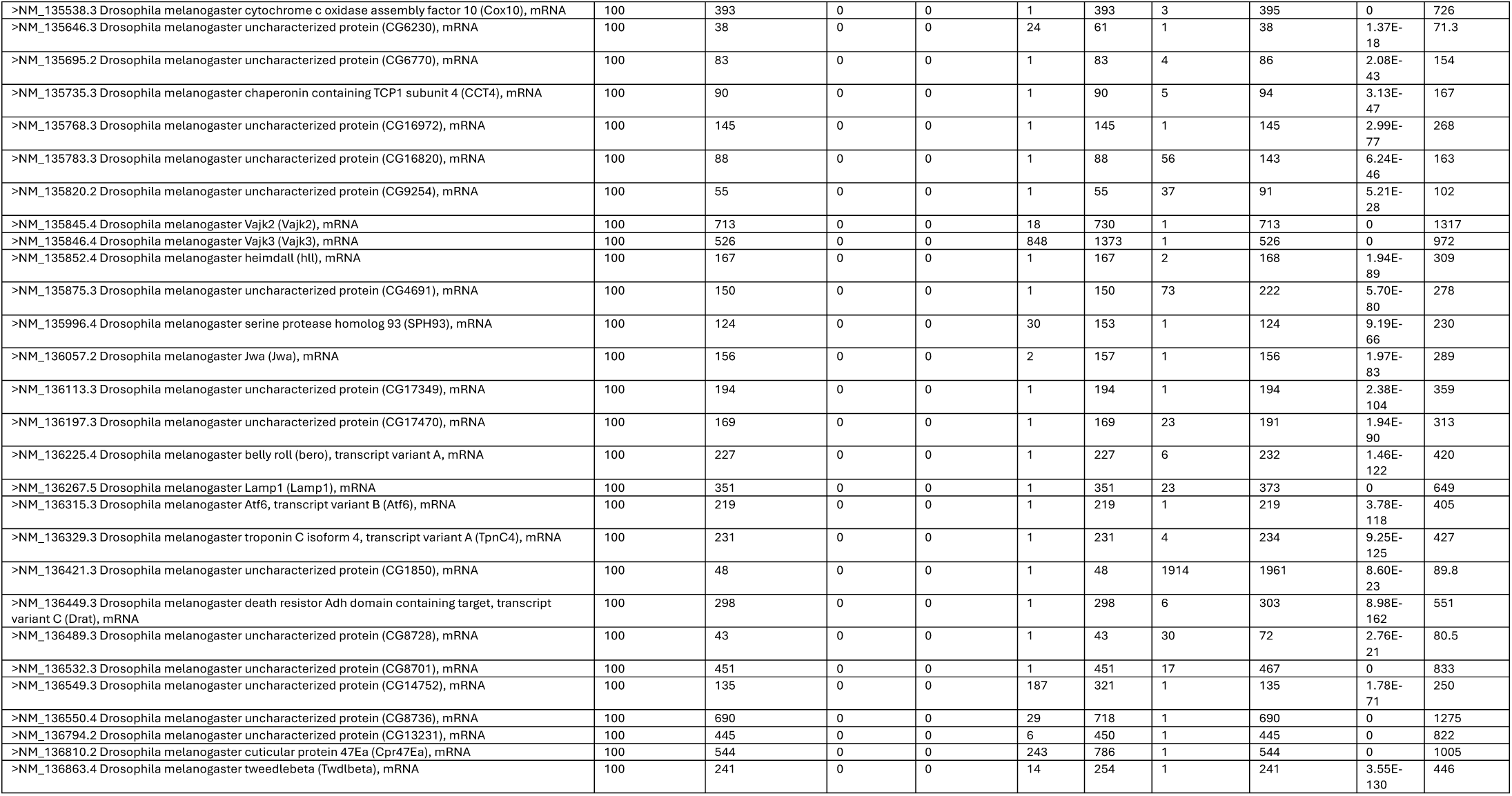
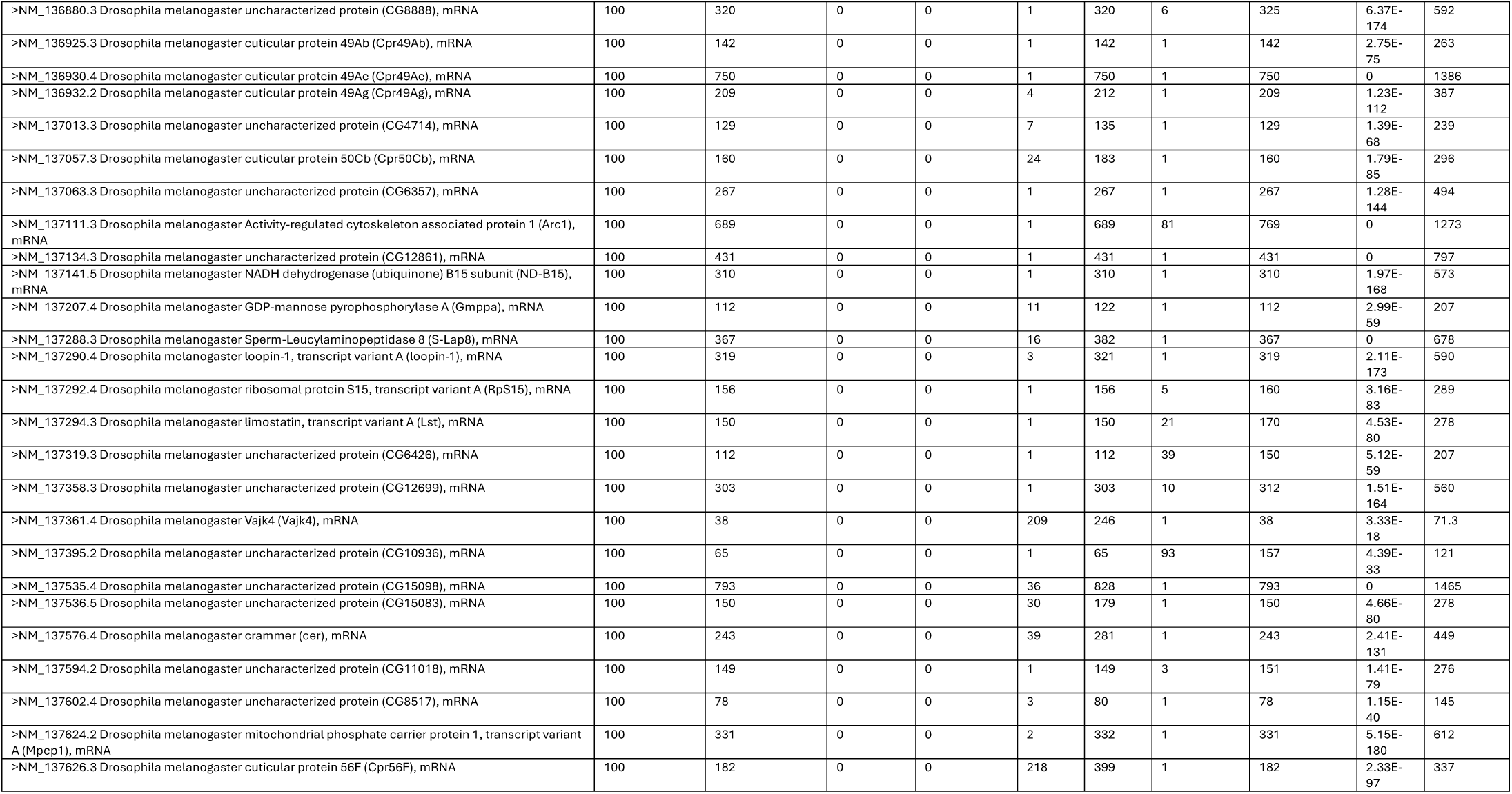
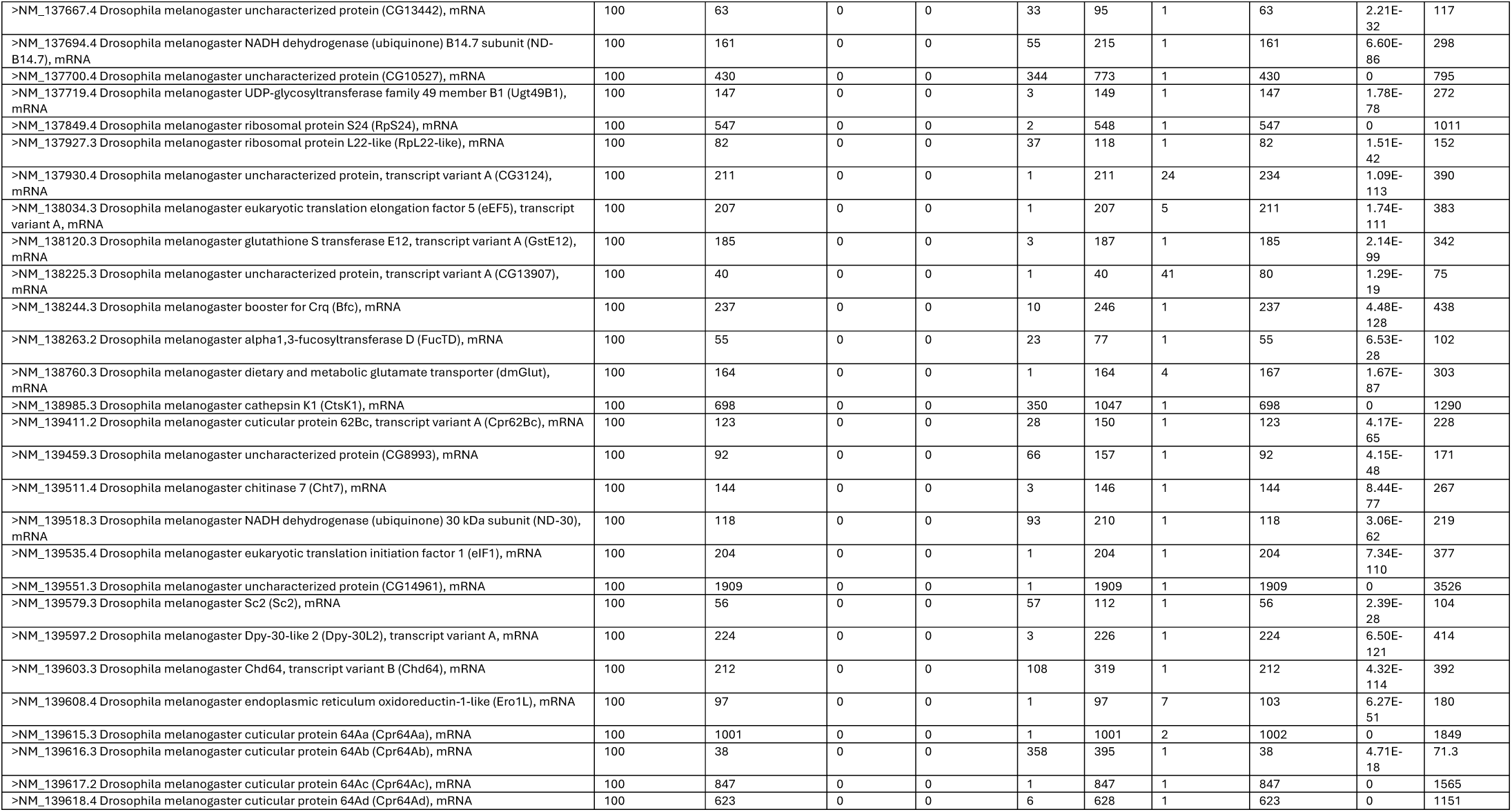
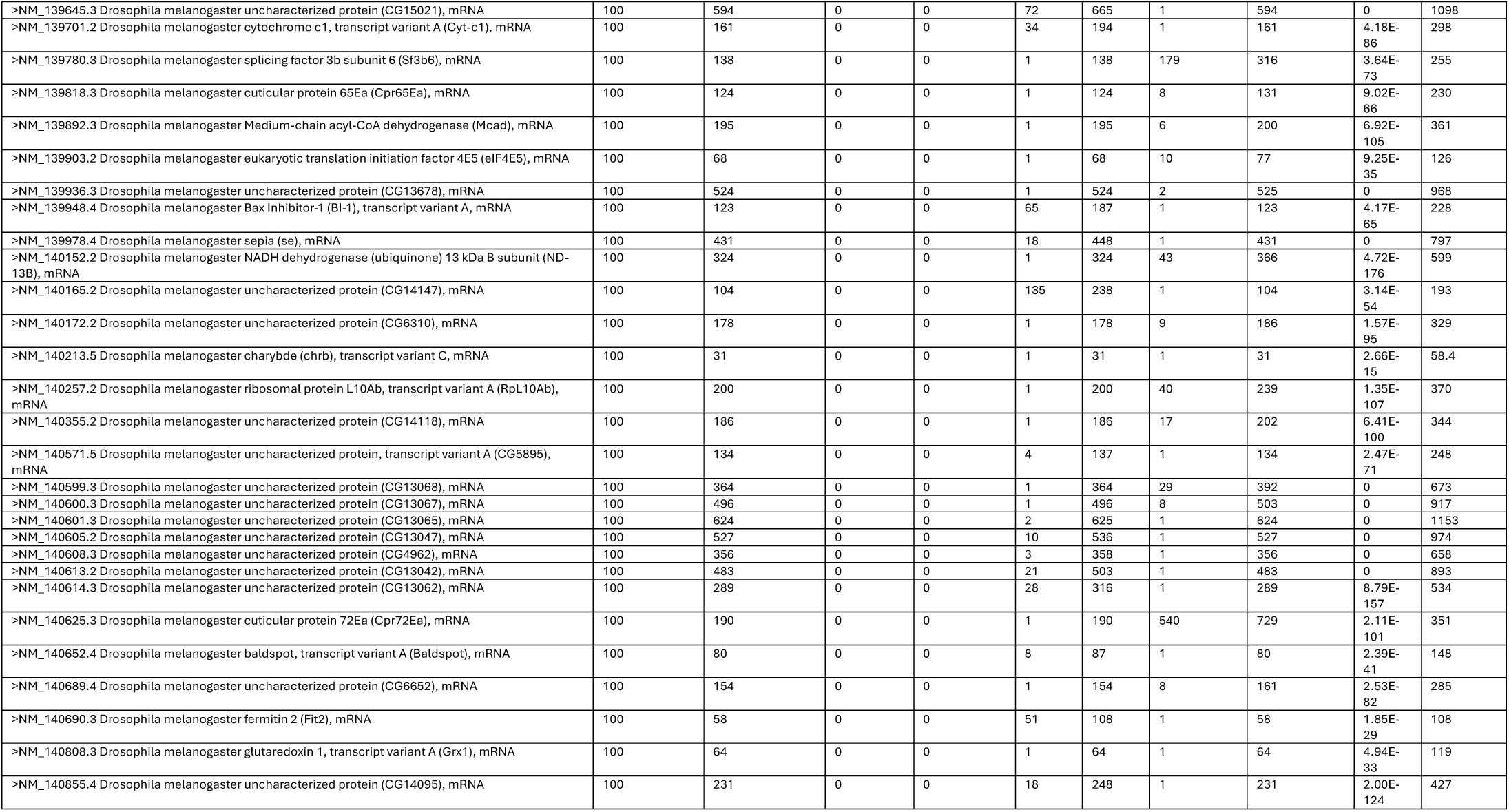
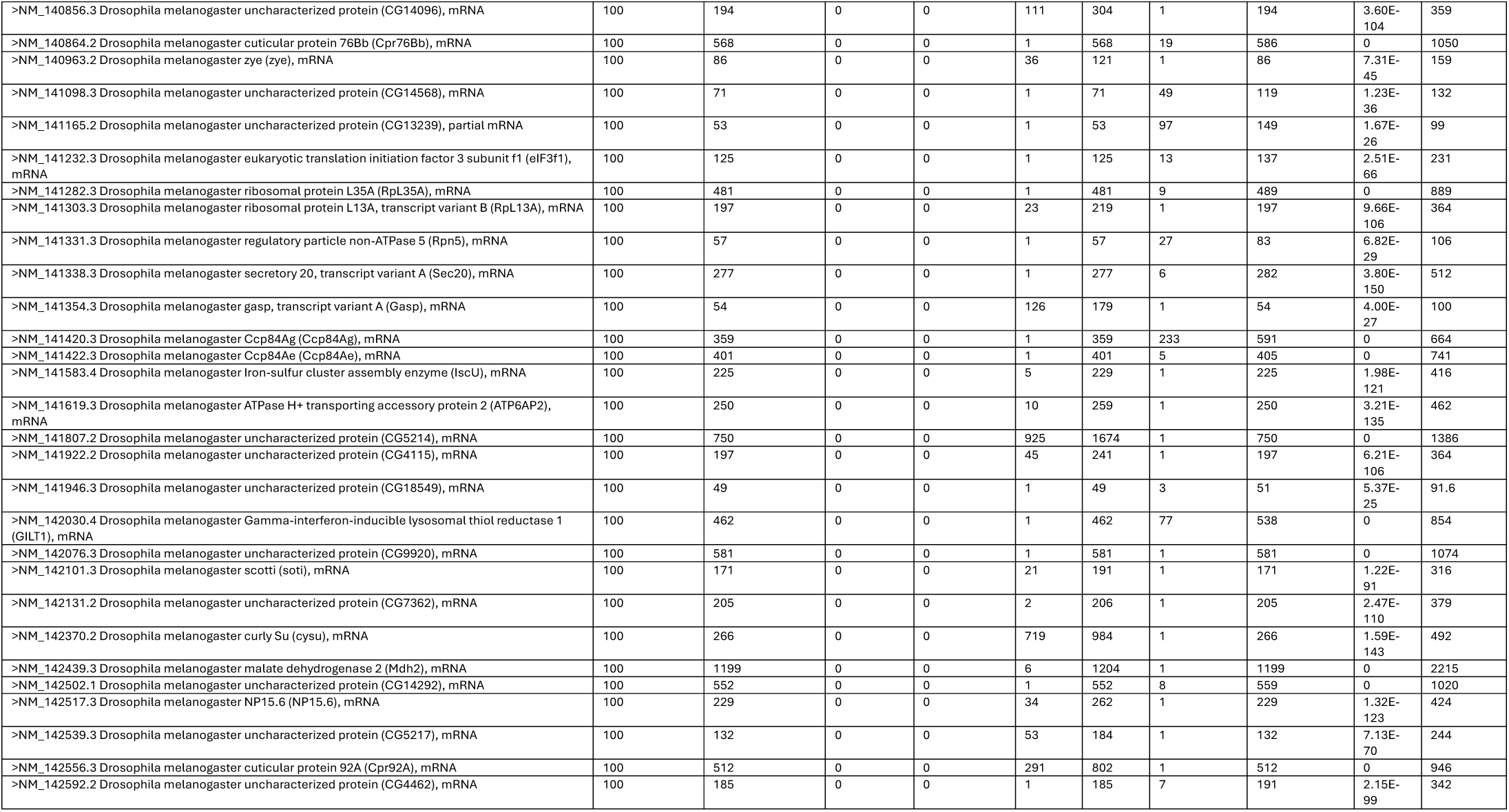
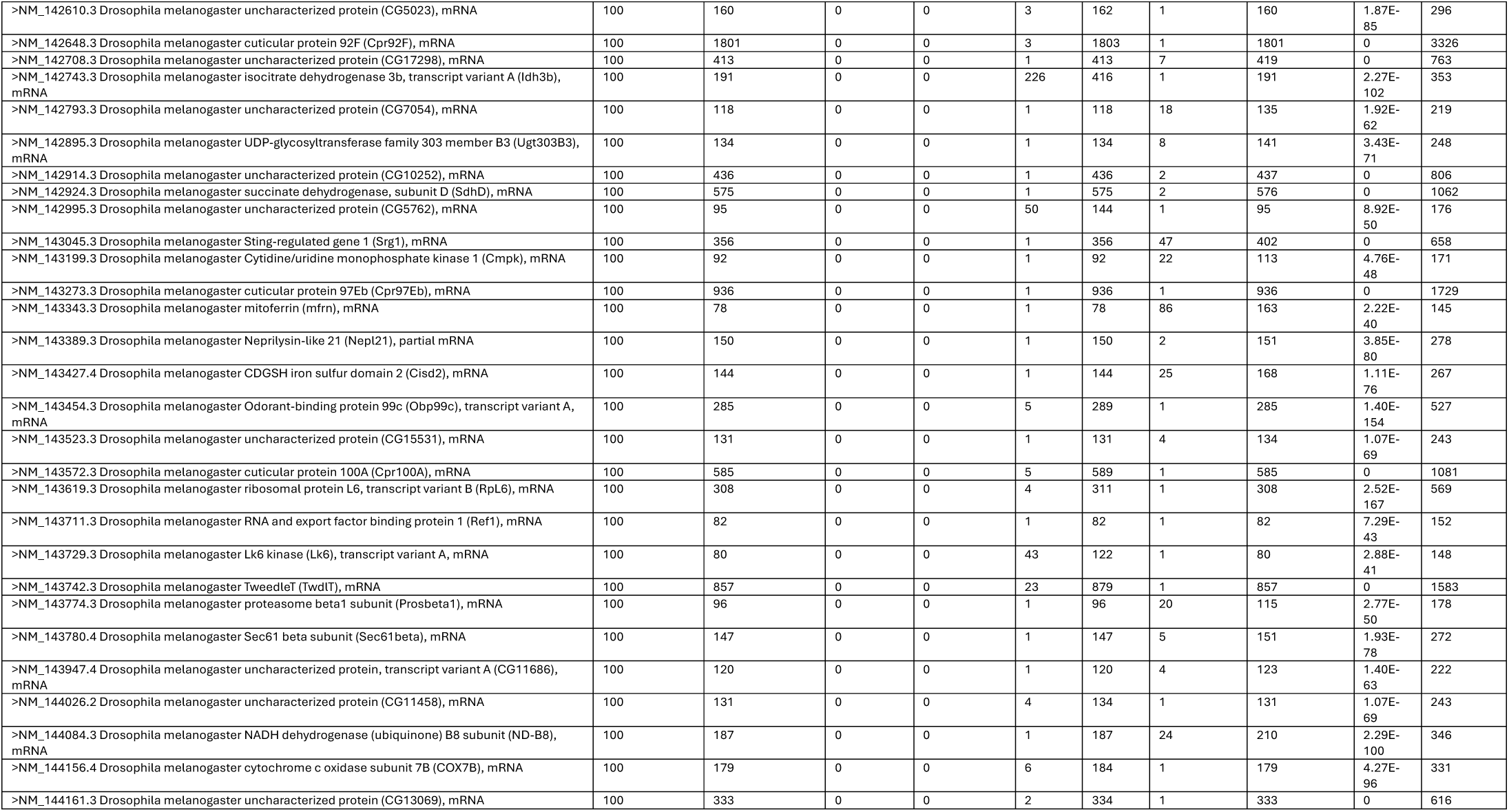
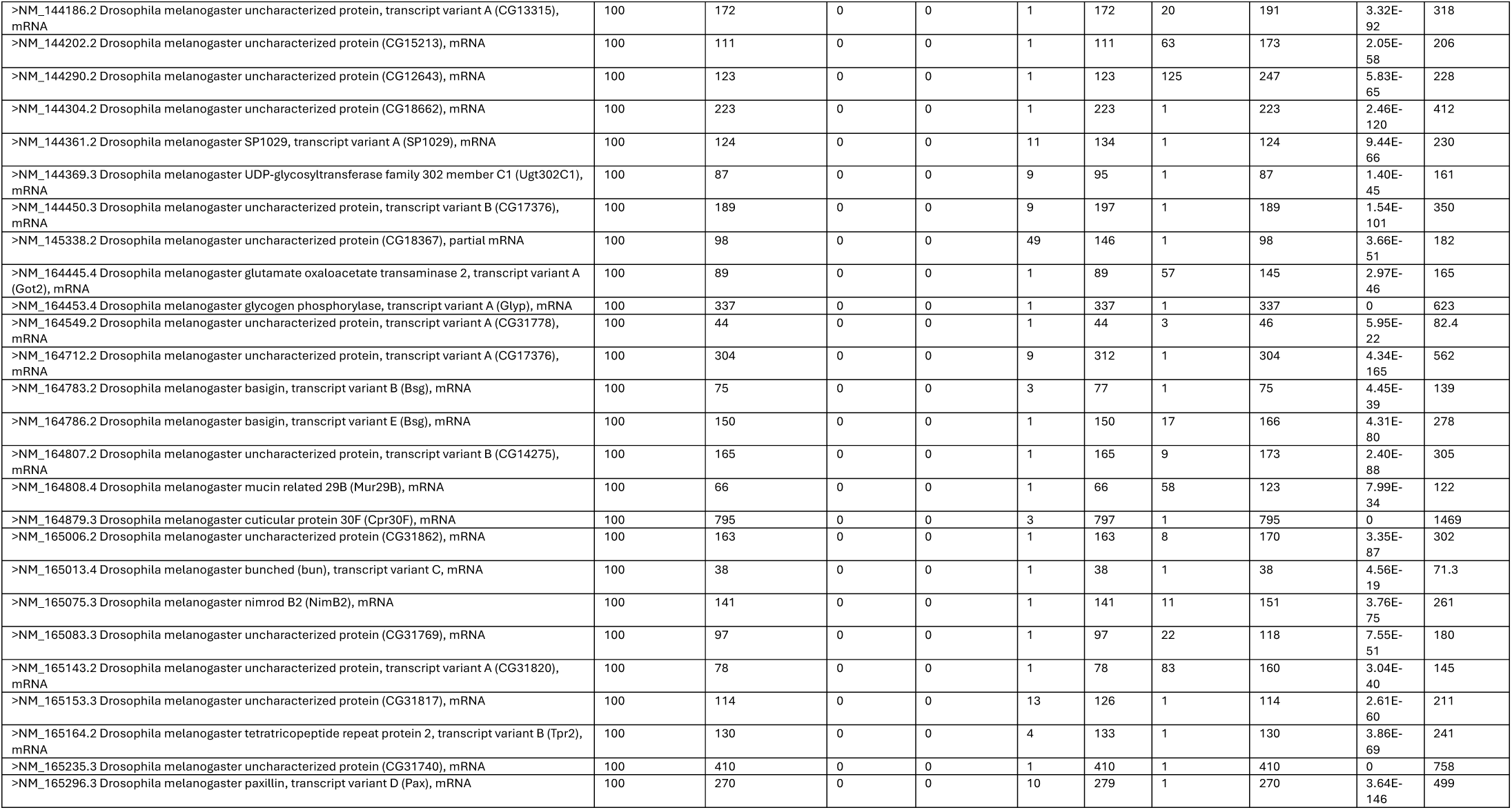
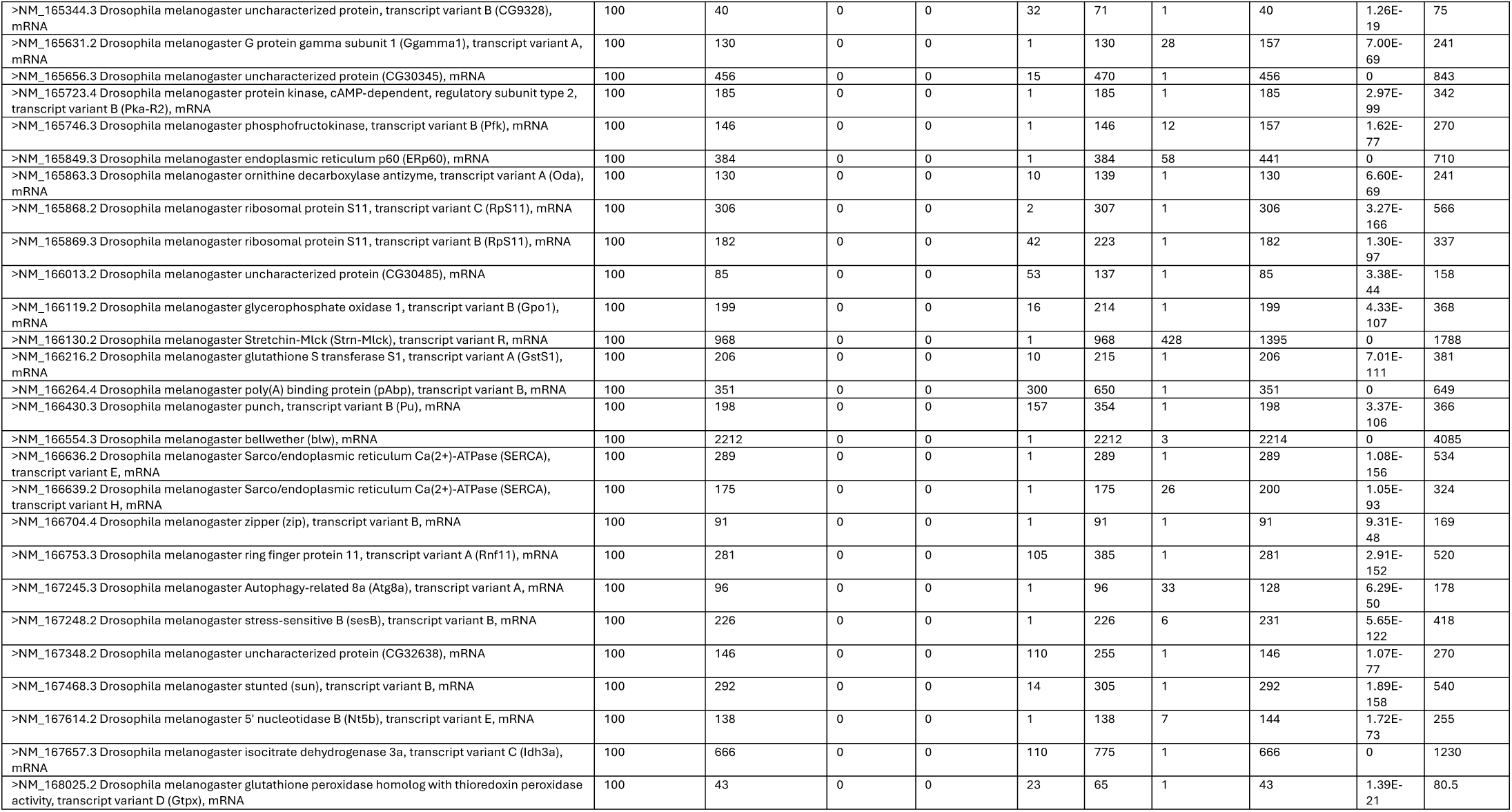
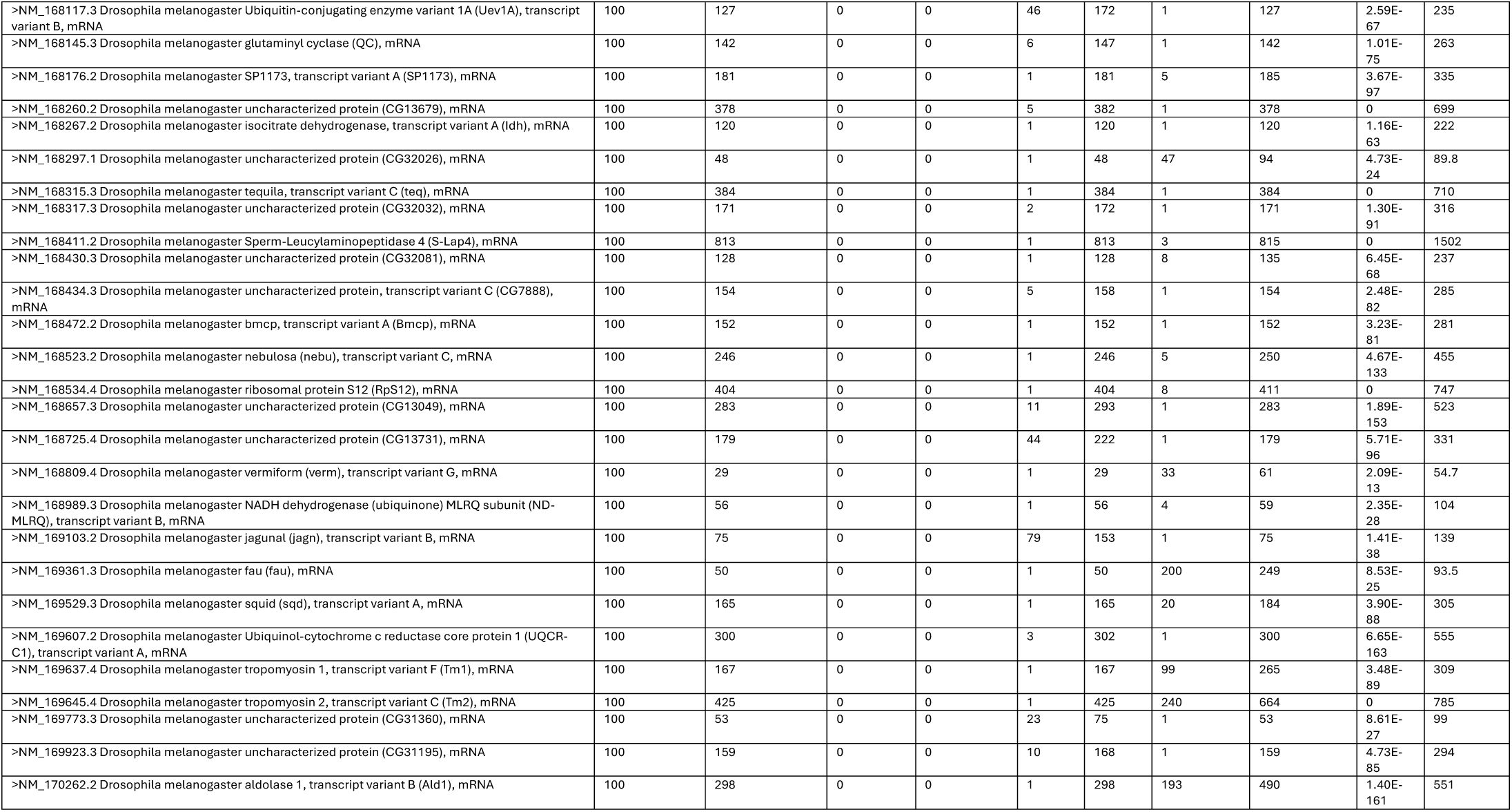
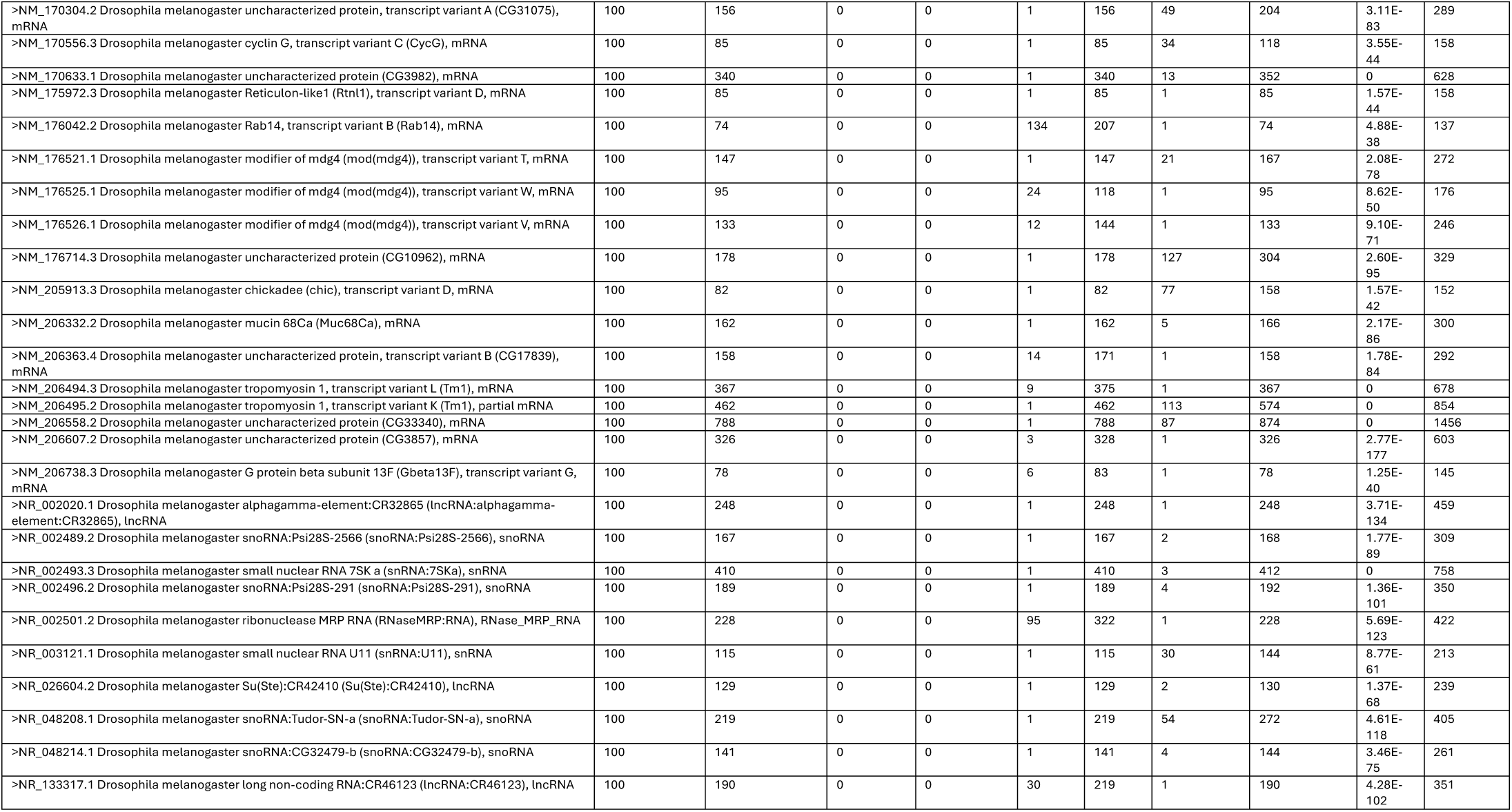
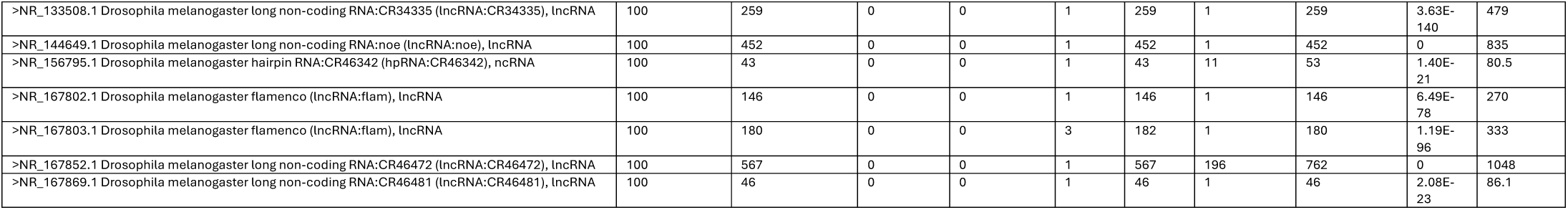
Genetic comparison between CV- and CV+ lines.

**S2 Table:**
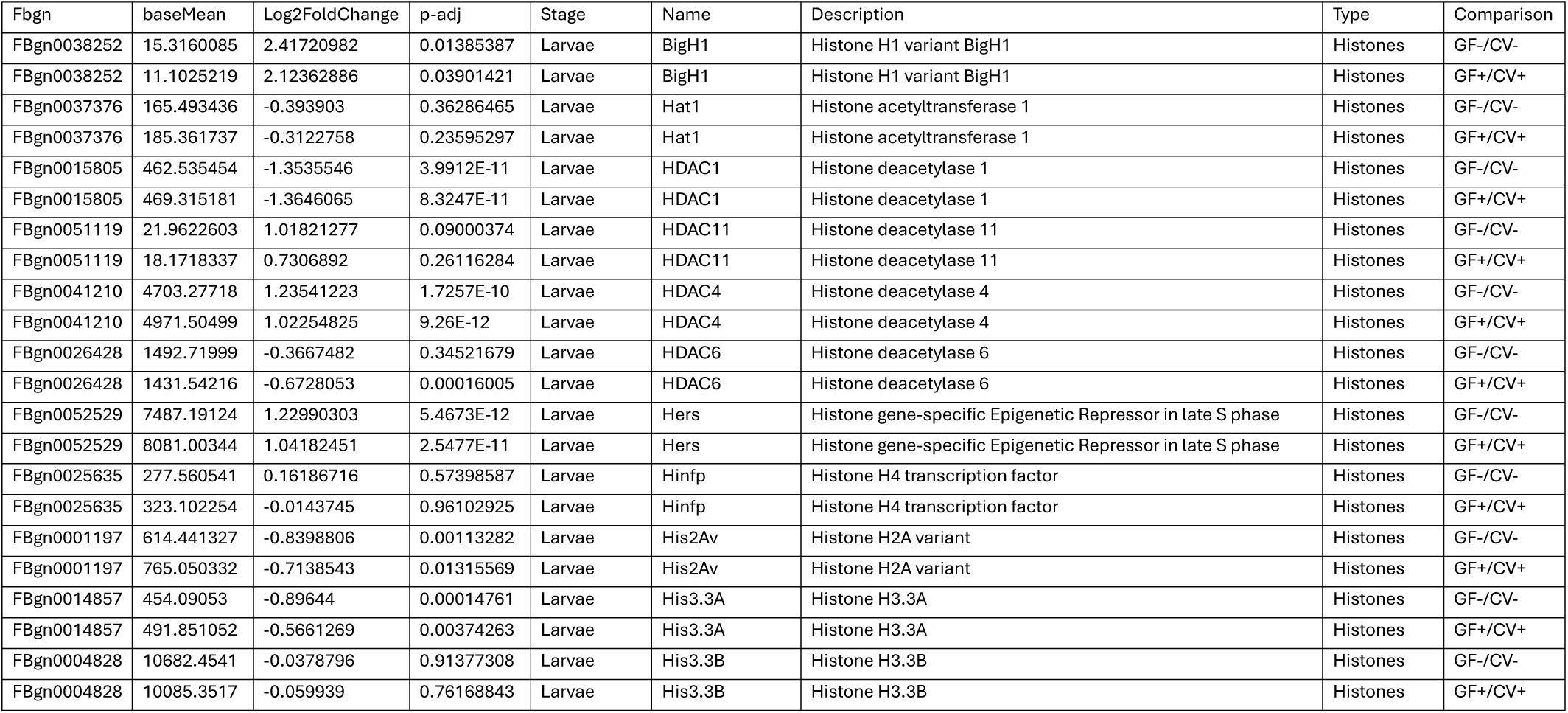
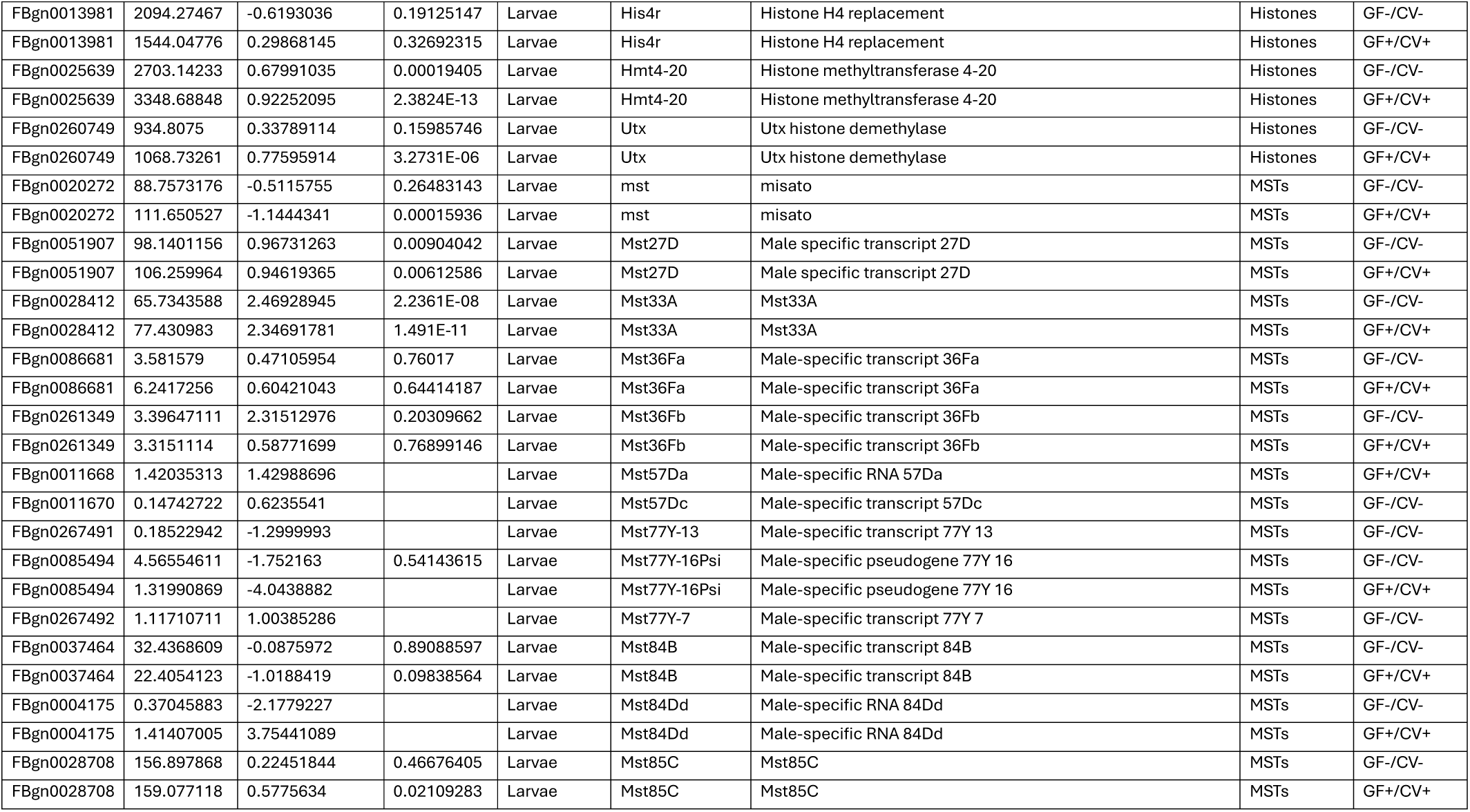
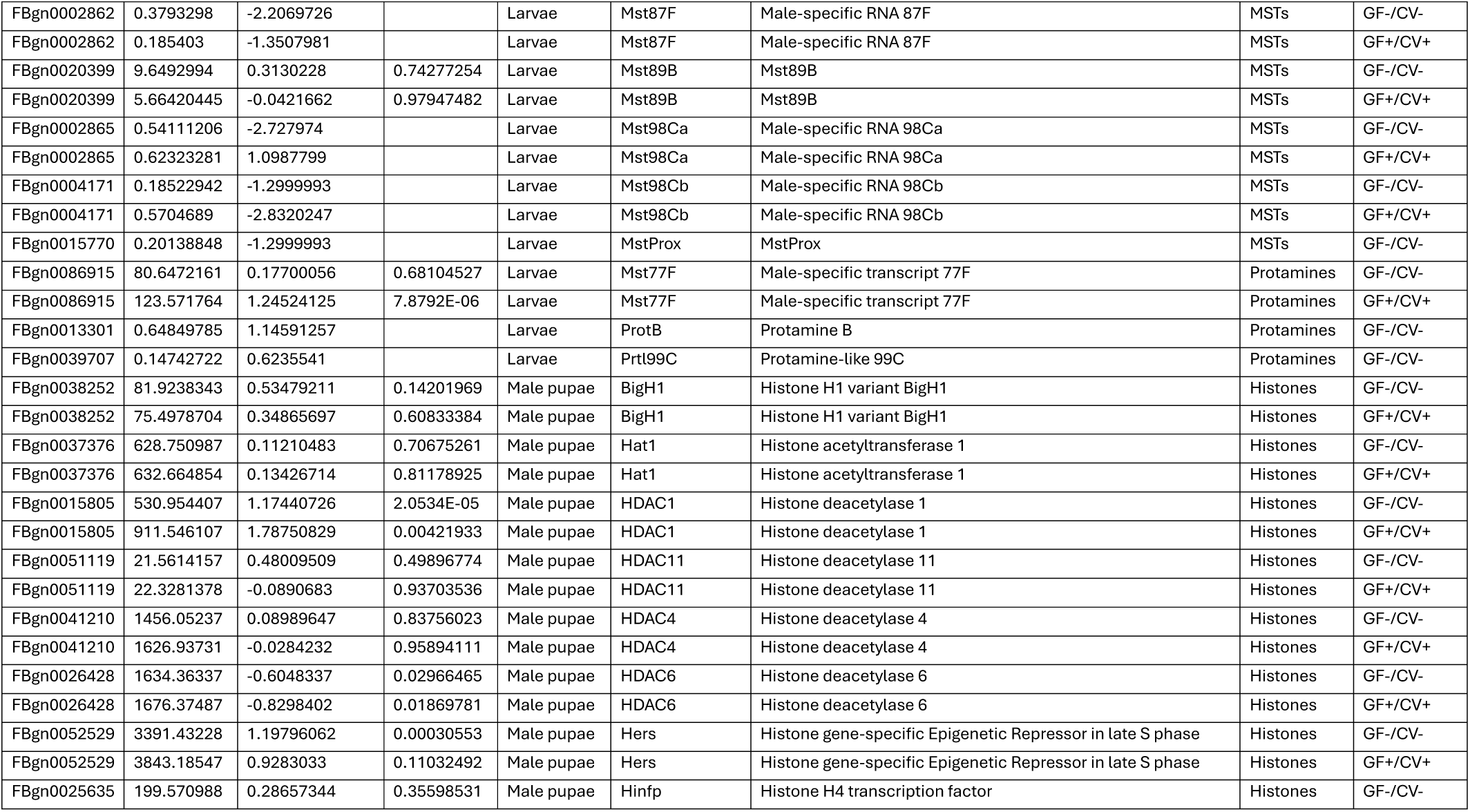
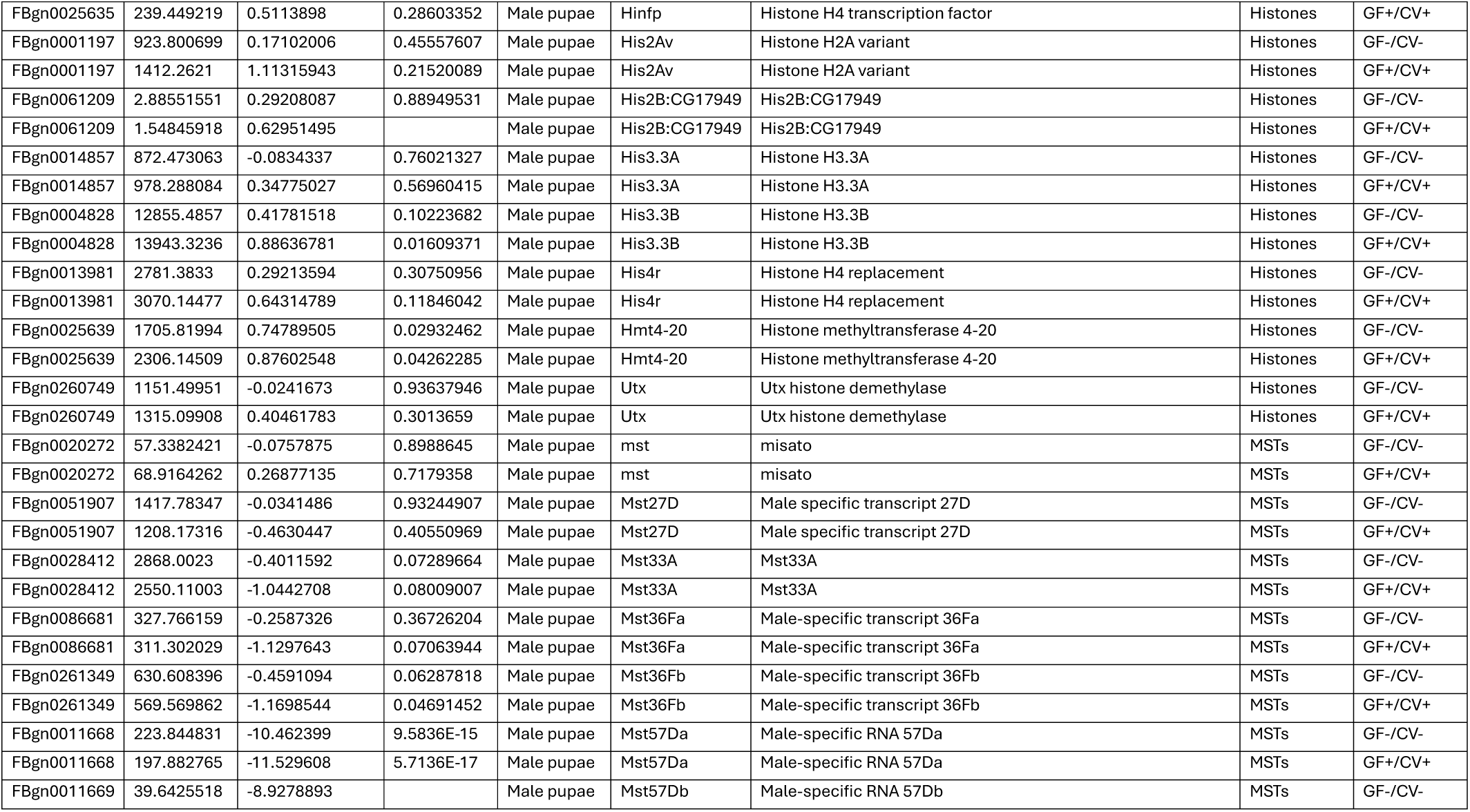
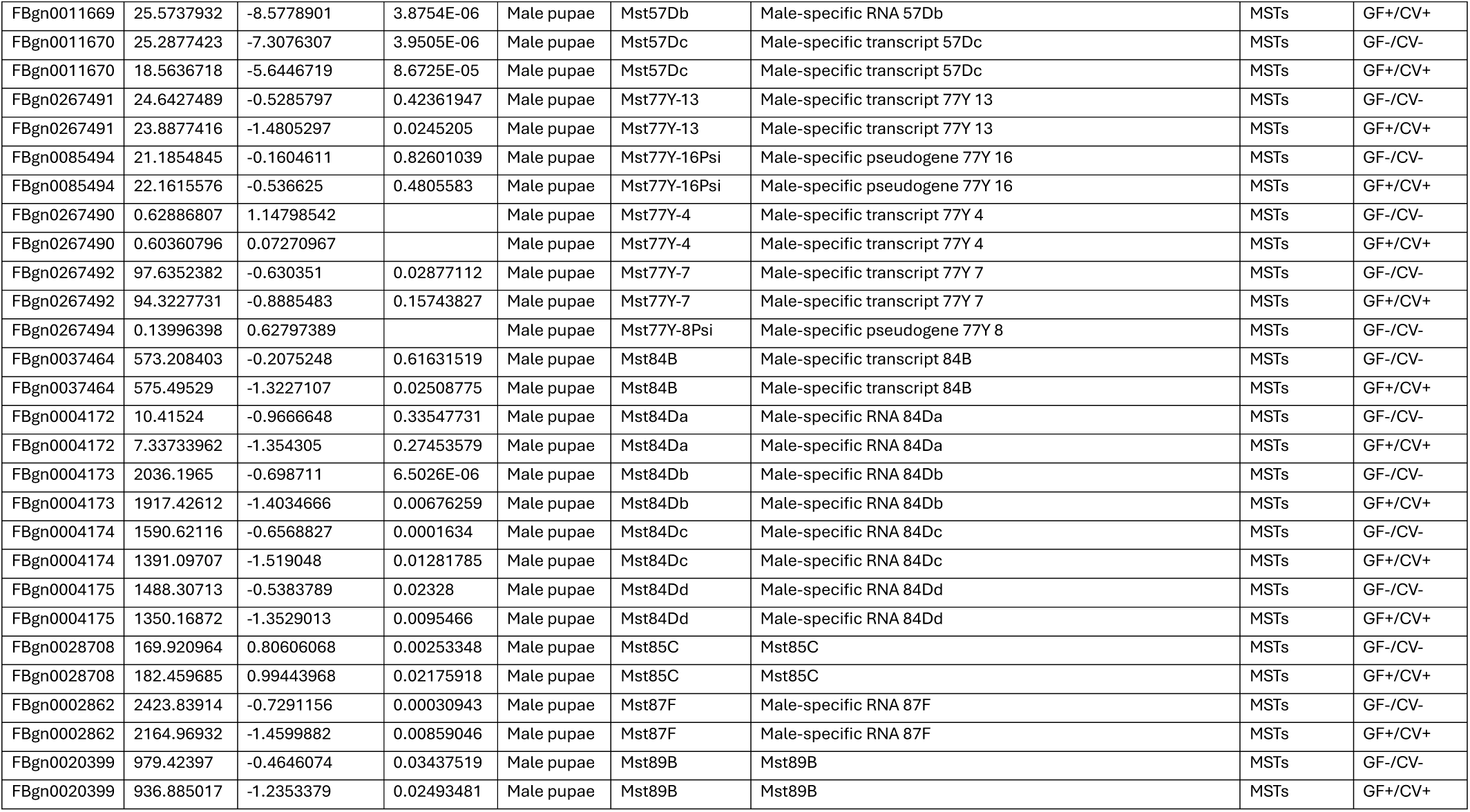
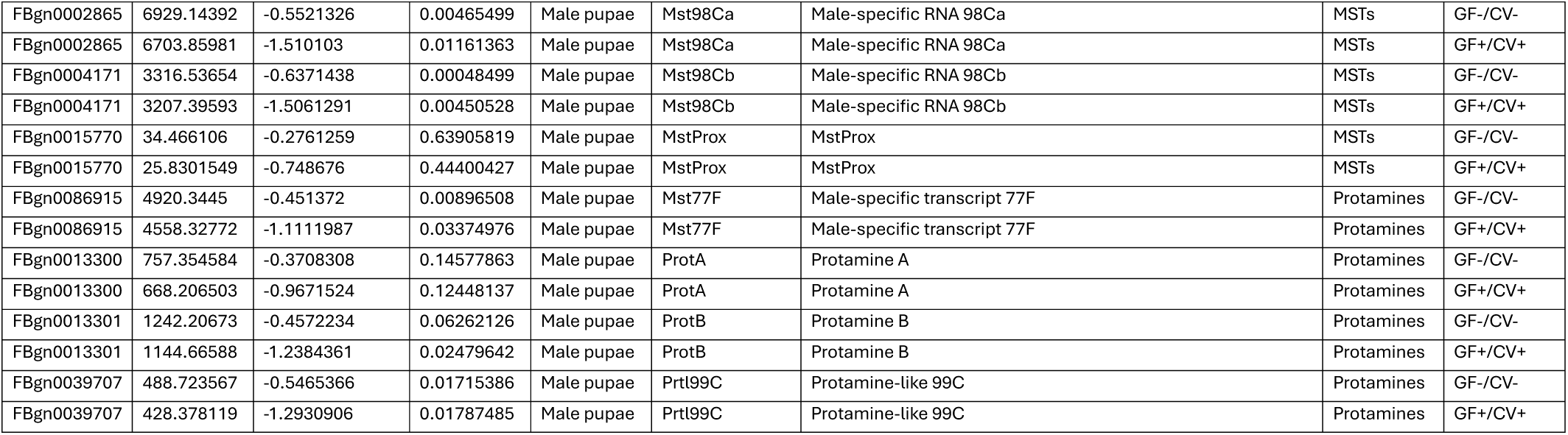
Differentially-expressed histone, protamine, and MST transcripts in GF+/CV+ and GF-/CV- larvae and male pupae.

**S3 Table:**
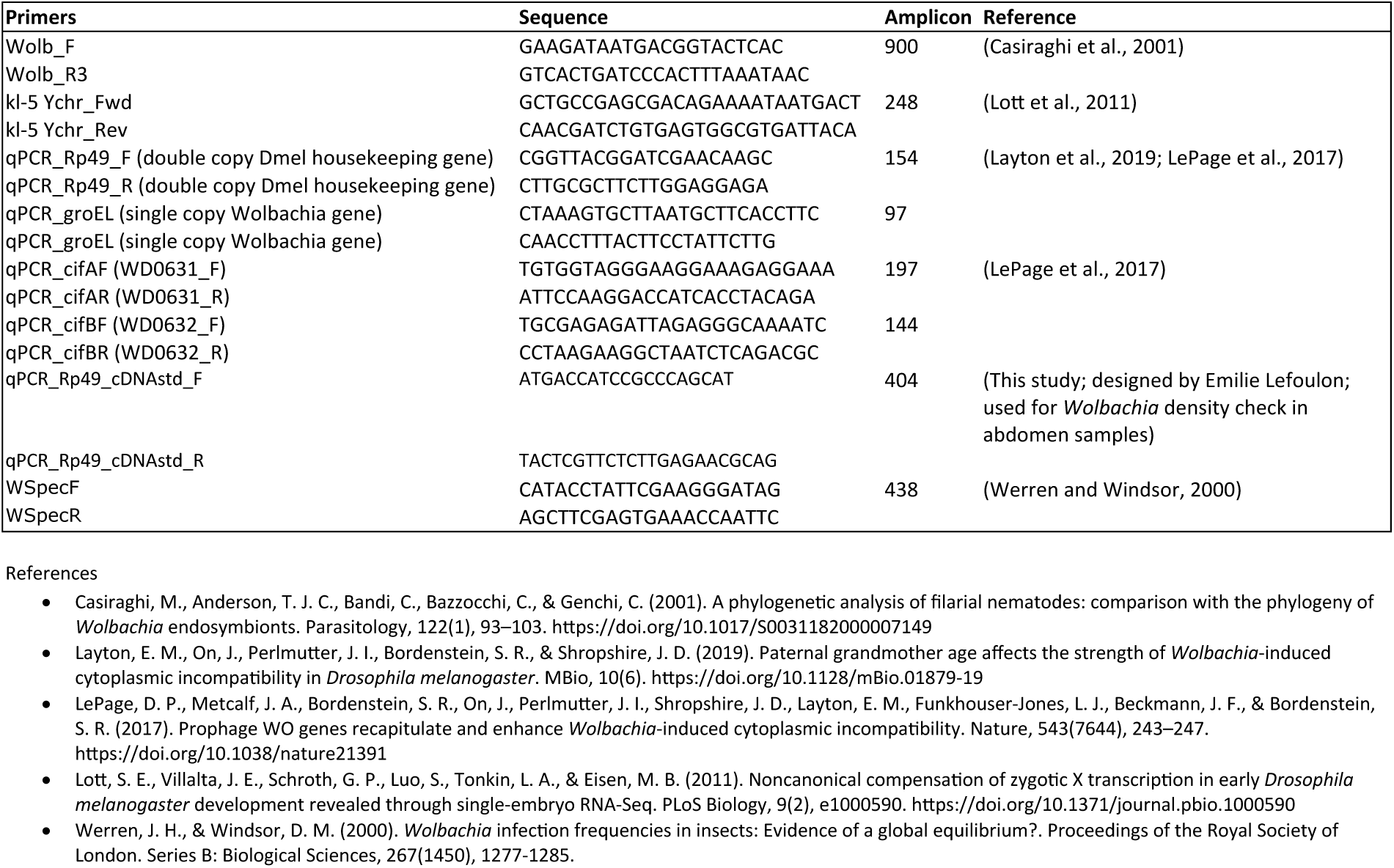
Primers used for measuring *Wolbachia* density, sex of pupae, and *cifA* and *cifB* expression.

## References

1. McFall-Ngai, M., Hadfield, M. G., Bosch, T. C. G., Carey, H. V., Domazet-Lošo, T., Douglas, A. E., Dubilier, N., Eberl, G., Fukami, T., Gilbert, S. F., Hentschel, U., King, N., Kjelleberg, S., Knoll, A. H., Kremer, N., Mazmanian, S. K., Metcalf, J. L., Nealson, K., Pierce, N. E., Rawls, J. F., Reid, A., Ruby, E. G., Rumpho, M., Sanders, J. G., Tautz, D. & Wernegreen, J. J. Animals in a bacterial world, a new imperative for the life sciences. Proceedings of the National Academy of Sciences 110, 3229–3236 (2013).

2. Bordenstein, S. R. & Theis, K. R. Host biology in light of the microbiome: Ten principles of holobionts and hologenomes. PLoS Biol. 13, e1002226 (2015).

3. Bordenstein, S. R., The Holobiont Biology Network (Gilbert, M. T. P., Ginnan, N., Malacrinò, A., Martino, M. E., Bahrndorff, S., Mundra, S., Martin, M. D., Theis, K. R., Hird, S. M., Caro-Quintero, A., Sharpton, T. J., Kohl, K. D., Barnes, C. J., Eisenhofer, R., Aizpurua, O., Andersen, S. B., Brealey, J. C., Noer, C. L., Medina, M., Limborg, M. T. & Alberdi, A). The disciplinary matrix of holobiont biology. Science (1979). 386, 731–732 (2024).

4. Brinker, P., Fontaine, M. C., Beukeboom, L. W. & Falcao Salles, J. Host, symbionts, and the microbiome: The missing tripartite interaction. Trends Microbiol. 27, 480–488 (2019).

5. Masson, F. & Lemaitre, B. Growing Ungrowable Bacteria: Overview and Perspectives on Insect Symbiont Culturability. Microbiol. Mol. Biol. Rev. 84, (2020).

6. Weinert, L. A., Araujo-Jnr, E. V., Ahmed, M. Z. & Welch, J. J. The incidence of bacterial endosymbionts in terrestrial arthropods. Proceedings of the Royal Society B: Biological Sciences 282, 20150249 (2015).

7. Kaur, R., Leigh, B. A., Ritchie, I. T. & Bordenstein, S. R. The Cif proteins from *Wolbachia* prophage WO modify sperm genome integrity to establish cytoplasmic incompatibility. PLoS Biol. 20, e3001584 (2022).

8. Shropshire, J. D., Leigh, B. & Bordenstein, S. R. Symbiont-mediated cytoplasmic incompatibility: What have we learned in 50 years? Elife 9, (2020).

9. Hoffmann, A. A., Turelli, M. & Harshman, L. G. Factors affecting the distribution of cytoplasmic incompatibility in *Drosophila simulans*. Genetics 126, 933–948 (1990).

10. Shropshire, J. D., Hamant, E., Conner, W. R. & Cooper, B. S. cifB-transcript levels largely explain cytoplasmic incompatibility variation across divergent Wolbachia. PNAS Nexus 1, (2022).

11. Landmann, F., Orsi, G. A., Loppin, B. & Sullivan, W. *Wolbachia*-mediated cytoplasmic incompatibility is associated with impaired histone deposition in the male pronucleus. PLoS Pathog. 5, e1000343 (2009).

12. Perez, C., Porter, J., Warecki, B. & Sullivan, W. *Wolbachia*-induced cytoplasmic incompatibility drives epigenetic and maternally-influenced post-embryonic defects. PLoS Pathog. 22, e1014180 (2026).

13. Ludington, W. B. & Ja, W. W. *Drosophila* as a model for the gut microbiome. PLoS Pathog. 16, e1008398 (2020).

14. Porter, J. & Sullivan, W. The cellular lives of *Wolbachia*. Nat. Rev. Microbiol. 21, 750–766 (2023).

15. Simhadri, R. K., Fast, E. M., Guo, R., Schultz, M. J., Vaisman, N., Ortiz, L., Bybee, J., Slatko, B. E. & Frydman, H. M. The gut commensal microbiome of Drosophila melanogaster is modified by the endosymbiont Wolbachia. mSphere 2, (2017).

16. Ye, Y. H., Seleznev, A., Flores, H. A., Woolfit, M. & McGraw, E. A. Gut microbiota in *Drosophila melanogaster* interacts with *Wolbachia* but does not contribute to *Wolbachia* - mediated antiviral protection. J. Invertebr. Pathol. 143, 18–25 (2017).

17. Blackie, L., Gaspar, P., Mosleh, S., Lushchak, O., Kong, L., Jin, Y., Zielinska, A. P., Cao, B., Mineo, A., Silva, B., Ameku, T., Lim, S. E., Mao, Y., Prieto-Godino, L., Schoborg, T., Varela, M., Mahadevan, L. & Miguel-Aliaga, I. The sex of organ geometry. Nature 630, 392–400 (2024).

18. Wolstenholme, D. R. A DNA and RNA-containing cytoplasmic body in *Drosophila melanogaster* and its relation to flies. Genetics 52, 949–975 (1965).

19. Bakula, M. A. The ecogenetics of a Drosophila-bacteria association. (1967).

20. Hoffmann, A. A., Hercus, M. & Dagher, H. Population dynamics of the *Wolbachia* infection causing cytoplasmic incompatibility in *Drosophila melanogaster*. Genetics 148, 221–231 (1998).

21. Blum, J. E., Fischer, C. N., Miles, J. & Handelsman, J. Frequent replenishment sustains the beneficial microbiome of *Drosophila melanogaster*. mBio 4, (2013).

22. Guerrero-Ferreira, R. C. & Nishiguchi, M. K. Differential gene expression in bacterial symbionts from loliginid squids demonstrates variation between mutualistic and environmental niches. Environ. Microbiol. Rep. 2, 514–523 (2010).

23. Wier, A. M., Nyholm, S. V., Mandel, M. J., Massengo-Tiassé, R. P., Schaefer, A. L., Koroleva, I., Splinter-BonDurant, S., Brown, B., Manzella, L., Snir, E., Almabrazi, H., Scheetz, T. E., Bonaldo, M. de F., Casavant, T. L., Soares, M. B., Cronan, J. E., Reed, J. L., Ruby, E. G. & McFall-Ngai, M. J. Transcriptional patterns in both host and bacterium underlie a daily rhythm of anatomical and metabolic change in a beneficial symbiosis. Proceedings of the National Academy of Sciences 107, 2259–2264 (2010).

24. Hansen, A. K. & Moran, N. A. Aphid genome expression reveals host–symbiont cooperation in the production of amino acids. Proceedings of the National Academy of Sciences 108, 2849–2854 (2011).

25. Mahan, M. J., Heithoff, D. M., Sinsheimer, R. L. & Low, D. A. Assessment of bacterial pathogenesis by analysis of gene expression in the host. Annu. Rev. Genet. 34, 139–164 (2000).

26. Lefranc, C., Fichant, A. & Storelli, G. Transient remodeling of gut metabolism supports juvenile growth and adult fitness in *Drosophila*. Nat. Commun. 17, 3458 (2026).

27. Storelli, G., Strigini, M., Grenier, T., Bozonnet, L., Schwarzer, M., Daniel, C., Matos, R. & Leulier, F. *Drosophila* perpetuates nutritional mutualism by promoting the fitness of its intestinal symbiont *Lactobacillus plantarum*. Cell Metab. 27, 362–377.e8 (2018).

28. Kaur, R., McGarry, A., Shropshire, J. D., Leigh, B. A. & Bordenstein, S. R. Prophage proteins alter long noncoding RNA and DNA of developing sperm to induce a paternal-effect lethality. Science (1979). 383, 1111–1117 (2024).

29. Rice, D. W., Sheehan, K. B. & Newton, I. L. G. Large-scale identification of *Wolbachia pipientis* effectors. Genome Biol. Evol. 9, 1925–1937 (2017).

30. Gutzwiller, F., Carmo, C. R., Miller, D. E., Rice, D. W., Newton, I. L. G., Hawley, R. S., Teixeira, L. & Bergman, C. M. Dynamics of *Wolbachia pipientis* gene expression across the *Drosophila melanogaster* Life Cycle. G3 Genes|Genomes|Genetics 5, 2843–2856 (2015).

31. Chung, M., Basting, P. J., Patkus, R. S., Grote, A., Luck, A. N., Ghedin, E., Slatko, B. E., Michalski, M., Foster, J. M., Bergman, C. M. & Hotopp, J. C. D. A Meta-analysis of *Wolbachia* transcriptomics reveals a stage-specific *Wolbachia* transcriptional response shared across different hosts. G3 Genes|Genomes|Genetics 10, 3243–3260 (2020).

32. Bordenstein, S. R. & Bordenstein, S. R. Eukaryotic association module in phage WO genomes from *Wolbachia*. Nat. Commun. 7, 13155 (2016).

33. LePage, D. P., Metcalf, J. A., Bordenstein, S. R., On, J., Perlmutter, J. I., Shropshire, J. D., Layton, E. M., Funkhouser-Jones, L. J., Beckmann, J. F. & Bordenstein, S. R. Prophage WO genes recapitulate and enhance *Wolbachia*-induced cytoplasmic incompatibility. Nature 543, 243–247 (2017).

34. Hamilton, W., Hardy, E., López-Madrigal, S., Phelps, M., Martin, M. & Newton, I. *Wolbachia* uses ankyrin repeats to target specific fly proteins. mBio 17, (2026).

35. Perlmutter, J. I., Meyers, J. E. & Bordenstein, S. R. Transgenic testing does not support a role for additional candidate genes in Wolbachia male killing or cytoplasmic incompatibility. mSystems 5, (2020).

36. Perlmutter, J. I. & Bordenstein, S. R. Microorganisms in the reproductive tissues of arthropods. Nat. Rev. Microbiol. 18, 97–111 (2020).

37. Yamada, R., Iturbe-Ormaetxe, I., Brownlie, J. C. & O’Neill, S. L. Functional test of the influence of *Wolbachia* genes on cytoplasmic incompatibility expression in *Drosophila melanogaster*. Insect Mol. Biol. 20, 75–85 (2011).

38. Kaur, R., Meier, C. J., McGraw, E. A., Hillyer, J. F. & Bordenstein, S. R. The mechanism of cytoplasmic incompatibility is conserved in *Wolbachia*-infected *Aedes aegypti* mosquitoes deployed for arbovirus control. PLoS Biol. 22, e3002573 (2024).

39. Ingham, P. The development of *Drosophila melanogaster*. Trends in Genetics 10, 299 (1994).

40. Williamson, A. & Lehmann, R. Germ cell development in *Drosophila*. Annu. Rev. Cell Dev. Biol. 12, 365–391 (1996).

41. Rathke, C., Baarends, W. M., Jayaramaiah-Raja, S., Bartkuhn, M., Renkawitz, R. & Renkawitz-Pohl, R. Transition from a nucleosome-based to a protamine-based chromatin configuration during spermiogenesis in *Drosophila*. J. Cell Sci. 120, 1689–1700 (2007).

42. Kornberg, R. D. & Lorch, Y. Twenty-five years of the nucleosome, fundamental particle of the eukaryote chromosome. Cell vol. 98 Preprint at 10.1016/S0092-8674(00)81958-3 (1999).

43. Jayaramaiah Raja, S. & Renkawitz-Pohl, R. Replacement by *Drosophila melanogaster* protamines and Mst77f of histones during chromatin condensation in late spermatids and role of sesame in the removal of these proteins from the male pronucleus. Mol. Cell. Biol. 25, 6165–6177 (2005).

44. Eren-Ghiani, Z., Rathke, C., Theofel, I. & Renkawitz-Pohl, R. Prtl99C Acts Together with Protamines and Safeguards Male Fertility in Drosophila. Cell Rep. 13, 2327–2335 (2015).

45. Kaur, R. & Bordenstein, S. R. Cytoplasmic incompatibility factor proteins from *Wolbachia* prophage are costly to sperm development in *Drosophila melanogaster*. Proceedings of the Royal Society B: Biological Sciences 292, (2025).

46. Ikeda, T., Ishikawa, H. & Sasaki, T. Infection density of *Wolbachia* and level of cytoplasmic incompatibility in the Mediterranean flour moth, *Ephestia kuehniella*. J. Invertebr. Pathol. 84, 1–5 (2003).

47. Ritchie, I. T., Needles, K. T., Leigh, B. A., Kaur, R. & Bordenstein, S. R. Transgenic cytoplasmic incompatibility persists across age and temperature variation in *Drosophila melanogaster*. iScience 25, 105327 (2022).

48. Bordenstein, S. R., Uy, J. J. & Werren, J. H. Host genotype determines cytoplasmic incompatibility type in the haplodiploid genus *Nasonia*. Genetics 164, 223–233 (2003).

49. Aranda-Díaz, A., Obadia, B., Dodge, R., Thomsen, T., Hallberg, Z. F., Güvener, Z. T., Ludington, W. B. & Huang, K. C. Bacterial interspecies interactions modulate pH-mediated antibiotic tolerance. Elife 9, (2020).

50. Storelli, G., Defaye, A., Erkosar, B., Hols, P., Royet, J. & Leulier, F. *Lactobacillus plantarum* promotes *Drosophila* systemic growth by modulating hormonal signals through TOR-dependent nutrient sensing. Cell Metab. 14, 403–414 (2011).

51. Newell, P. D. & Douglas, A. E. Interspecies Interactions determine the impact of the gut microbiota on nutrient allocation in *Drosophila melanogaster*. Appl. Environ. Microbiol. 80, 788–796 (2014).

52. Sommer, A. J. & Newell, P. D. Metabolic basis for mutualism between gut bacteria and its impact on the *Drosophila melanogaster* Host. Appl. Environ. Microbiol. 85, (2019).

53. Nunes, R. D. & Drummond-Barbosa, D. A high-sugar diet, but not obesity, reduces female fertility in *Drosophila melanogaster*. Development 150, (2023).

54. Horváth, B. & Kalinka, A. T. Effects of larval crowding on quantitative variation for development time and viability in *Drosophila melanogaster*. Ecol. Evol. 6, 8460–8473 (2016).

55. Heys, C., Lizé, A., Blow, F., White, L., Darby, A. & Lewis, Z. J. The effect of gut microbiota elimination in *Drosophila melanogaster*: A how-to guide for host–microbiota studies. Ecol. Evol. 8, 4150–4161 (2018).

56. Ridley, E. V., Wong, A. C.-N., Westmiller, S. & Douglas, A. E. Impact of the resident microbiota on the nutritional phenotype of *Drosophila melanogaster*. PLoS One 7, e36765 (2012).

57. Chaston, J. M., Dobson, A. J., Newell, P. D. & Douglas, A. E. Host genetic control of the microbiota mediates the *Drosophila* nutritional phenotype. Appl. Environ. Microbiol. 82, (2016).

58. Sherwin, E., Bordenstein, S. R., Quinn, J. L., Dinan, T. G. & Cryan, J. F. Microbiota and the social brain. Science (1979). 366, (2019).

59. Dang, A. T. & Marsland, B. J. Microbes, metabolites, and the gut–lung axis. Mucosal Immunol. 12, 843–850 (2019).

60. Argaw-Denboba, A., Schmidt, T. S. B., Di Giacomo, M., Ranjan, B., Devendran, S., Mastrorilli, E., Lloyd, C. T., Pugliese, D., Paribeni, V., Dabin, J., Pisaniello, A., Espinola, S., Crevenna, A., Ghosh, S., Humphreys, N., Boruc, O., Sarkies, P., Zimmermann, M., Bork, P. & Hackett, J. A. Paternal microbiome perturbations impact offspring fitness. Nature 629, 652–659 (2024).

61. Hong, C., Lalsiamthara, J., Ren, J., Sang, Y. & Aballay, A. Microbial colonization induces histone acetylation critical for inherited gut-germline-neural signaling. PLoS Biol. 19, e3001169 (2021).

62. Layton, E. M., On, J., Perlmutter, J. I., Bordenstein, S. R. & Shropshire, J. D. Paternal grandmother age affects the strength of *Wolbachia*-induced cytoplasmic incompatibility in *Drosophila melanogaster*. mBio 10, (2019).

63. Bourtzis, K., Nirgianaki, A., Markakis, G. & Savakis, C. *Wolbachia* infection and cytoplasmic incompatibility in *Drosophila species*. Genetics 144, 1063–1073 (1996).

64. Cooper, B. S., Ginsberg, P. S., Turelli, M. & Matute, D. R. *Wolbachia* in the *Drosophila yakuba* Complex: Pervasive frequency variation and weak cytoplasmic incompatibility, but no apparent effect on reproductive isolation. Genetics 205, 333–351 (2017).

65. Shropshire, J. D., On, J., Layton, E. M., Zhou, H. & Bordenstein, S. R. One prophage WO gene rescues cytoplasmic incompatibility in *Drosophila melanogaster*. Proceedings of the National Academy of Sciences 115, 4987–4991 (2018).

66. Reynolds, K. T. & Hoffmann, A. A. Male age, host effects and the weak expression or non-expression of cytoplasmic incompatibility in *Drosophila* strains infected by maternally transmitted *Wolbachia*. Genet. Res. 80, (2002).

67. Yamada, R., Floate, K. D., Riegler, M. & O’Neill, S. L. Male development time influences the strength of *Wolbachia*-induced cytoplasmic incompatibility expression in *Drosophila melanogaster*. Genetics 177, 801–808 (2007).

68. Numajiri, Y., Kondo, N. I., Toquenaga, Y. & Kageyama, D. Intraspecies variation in cytoplasmic incompatibility intensity in the bean beetle *Callosobruchus analis*. Evol. Ecol. 38, 861–870 (2024).

69. Sasaki, T., Kubo, T. & Ishikawa, H. Interspecific transfer of *Wolbachia* between two lepidopteran insects expressing cytoplasmic incompatibility: A *Wolbachia* variant naturally infecting *cadra cautella* causes male killing in *Ephestia kuehniella*. Genetics 162, 1313–1319 (2002).

70. Suwanchaisri, K., Roddee, J. & Wangkeeree, J. *Wolbachia* transinfection and effect on the biological traits of *Matsumuratettix hiroglyphicus* (Matsumura), the leafhopper vector of sugarcane white leaf disease. Agriculture 14, 1236 (2024).

71. Hughes, G. L., Dodson, B. L., Johnson, R. M., Murdock, C. C., Tsujimoto, H., Suzuki, Y., Patt, A. A., Cui, L., Nossa, C. W., Barry, R. M., Sakamoto, J. M., Hornett, E. A. & Rasgon, J. L. Native microbiome impedes vertical transmission of *Wolbachia* in *Anopheles* mosquitoes. Proceedings of the National Academy of Sciences 111, 12498–12503 (2014).

72. Walker, T., Johnson, P. H., Moreira, L. A., Iturbe-Ormaetxe, I., Frentiu, F. D., McMeniman, C. J., Leong, Y. S., Dong, Y., Axford, J., Kriesner, P., Lloyd, A. L., Ritchie, S. A., O’Neill, S. L. & Hoffmann, A. A. The *w*Mel *Wolbachia* strain blocks dengue and invades caged *Aedes aegypti* populations. Nature 476, (2011).

73. Blagrove, M. S. C., Arias-Goeta, C., Failloux, A. B. & Sinkins, S. P. *Wolbachia* strain *w*Mel induces cytoplasmic incompatibility and blocks dengue transmission in *Aedes albopictus*. Proc. Natl. Acad. Sci. U. S. A. 109, (2012).

74. O’Neill, S. L. The use of *Wolbachia* by the world mosquito program to interrupt transmission of *Aedes aegypti* transmitted viruses. in Advances in Experimental Medicine and Biology vol. 1062 (2018).

75. Suito, T., Nagao, K., Juni, N., Hara, Y., Sokabe, T., Atomi, H. & Umeda, M. Regulation of thermoregulatory behavior by commensal bacteria in *Drosophila*. Biosci. Biotechnol. Biochem. 86, 1060–1070 (2022).

76. Koyle, M. L., Veloz, M., Judd, A. M., Wong, A. C.-N., Newell, P. D., Douglas, A. E. & Chaston, J. M. Rearing the fruit fly *Drosophila melanogaster* under axenic and gnotobiotic conditions. Journal of Visualized Experiments 10.3791/54219 (2016) doi:10.3791/54219.

77. Shropshire, J. D., Hamant, E. & Cooper, B. S. Male age and *Wolbachia* dynamics: investigating how fast and why bacterial densities and cytoplasmic incompatibility strengths vary. mBio 12, (2021).

78. Chandler, J. A., Innocent, L. V., Martinez, D. J., Huang, I. L., Yang, J. L., Eisen, M. B. & Ludington, W. B. Microbiome-by-ethanol interactions impact *Drosophila melanogaster* fitness, physiology, and behavior. iScience 25, 104000 (2022).

79. da Silva Soares, N. F., Quagliariello, A., Yigitturk, S. & Martino, M. E. Gut microbes predominantly act as living beneficial partners rather than raw nutrients. Sci. Rep. 13, 11981 (2023).

80. Lott, S. E., Villalta, J. E., Schroth, G. P., Luo, S., Tonkin, L. A. & Eisen, M. B. Noncanonical compensation of zygotic x transcription in early *Drosophila melanogaster* development revealed through single-embryo RNA-Seq. PLoS Biol. 9, e1000590 (2011).

